# Paralemmin-1 controls the nanoarchitecture of the neuronal submembrane cytoskeleton

**DOI:** 10.1101/2024.07.23.604764

**Authors:** Victor Macarrón-Palacios, Jasmine Hubrich, Maria Augusta do Rego Barros Fernandes Lima, Nicole G. Metzendorf, Simon Kneilmann, Marleen Trapp, Claudio Acuna, Annarita Patrizi, Elisa D’Este, Manfred W. Kilimann

## Abstract

The Membrane-associated Periodic Skeleton (MPS) is a specialized submembrane cytoskeleton of neuronal cells, characterized by a highly ordered 190 nm periodic lattice, with emerging functions in mechanical resilience, inter- and intracellular signaling, and action potential transmission. Here, we identify Paralemmin-1 (Palm1) as a new component and regulator of the MPS. Palm1 binds to the N-terminal region of βII-spectrin, a core MPS component, and is periodically organized along the axon in hippocampal neurons. Applying the 3D imaging power of MINFLUX, we locate Palm1 in close proximity (<20 nm) to the actin-capping protein and MPS component adducin. Functionally, Palm1 overexpression enhances the degree of periodicity of several MPS proteins (βII-spectrin, adducin, and ankyrinB) without altering their local concentrations, while the knock-out severely compromises the MPS structure and modifies electrophysiological properties of neurons. Both the MPS-binding and remodelling activities of Palm1 are abolished by mutating a single amino acid (W54A) in the conserved Paralemmin sequence motif. Our findings identify Palm1 as the first protein specifically dedicated to organizing the MPS, and will advance the understanding of the regulation of MPS assembly and remodelling, as well as of the Paralemmin protein family.

## Introduction

The Membrane-associated Periodic Skeleton (MPS) is a cytoskeletal structure underneath the plasma membrane of neurons and glial cells^1–3^. This lattice is most pronounced along axons, where actin filaments form ring-like structures, longitudinally separated by ∼190 nm intervals^2,4^. The spacing between the rings is maintained by spectrin tetramers, formed by two α/β-spectrin dimers interacting head-to-head near the C-terminus of β-spectrin. The N-terminus of β-spectrin binds actin, and this interaction is promoted by adducin^5,6^. In neurons, spectrin tetramers are composed of αII-spectrin and cell-compartment-specific isoforms of β-spectrin: βII-spectrin in axons and dendrites, βIII-spectrin in dendrites, and βIV-spectrin in the axon initial segment (AIS) and nodes of Ranvier^7–10^.

Docking proteins, such as ankyrins, bind to the MPS near the C-terminus of β-spectrin and act as adaptors for transmembrane proteins (e.g. ion channels, adhesion molecules, and receptors), which therefore also follow a ∼190 nm periodic organization^8,11^. Recent proteomic studies identified over 400 potential components of the MPS^12^, in line with the growing number of biological functions of this structure, which include mechanical support, and the mediation of intercellular (cell-cell interactions and action potential regulation) and intracellular (transport, signaling pathways) communication. However, the mechanisms regulating the MPS assembly and modulation are only starting to emerge. One factor that determines the quality of the periodicity is the concentration of βII-spectrin^9,13^. In axons, βII-spectrin is twice as abundant as in dendrites, where the MPS is less regular. Increasing the concentration of βII-spectrin in dendrites, by transient overexpression or as a consequence of ankyrinB (ankB) depletion, enhances long-range MPS periodicity in all neurites^13^. On the other hand, signaling proteins (G-protein-coupled receptors, cell-adhesion molecules, receptor tyrosine-kinases) can be recruited into the MPS upon activation, and induce a reversible calpain-mediated βII-spectrin degradation, which affects the architecture of the MPS^14^. A similar pathway is triggered by starvation and axonal damage^15,16^. However, no MPS component with a dedicated role in controlling its degree of periodicity has been identified.

Paralemmin-1 (Palm1) was discovered as a constituent of synaptic plasma membranes, attached to the cytoplasmic face of membranes by a prenyl-dipalmitoyl lipid anchor^17^. It is a member of the paralemmin family, which additionally comprises Paralemmin-2 (Palm2), Paralemmin-3 (Palm3), and Palmdelphin (Palmd)^18^. The overexpression of Palm1 in cell culture induces cell expansion and extension of processes, while in hippocampal neurons it enhances dendritic spine formation and synapse maturation, suggesting a role in cell shape control^17,19^. Interactions with the actin cytoskeleton have been identified for the paralemmin isoform Palmd. The lipid-anchored splice variant of Palmd co-immunoprecipitates with adducin, and enhances the branching of neuronal precursor cells^20^. The cytosolic splice variant of Palmd modulates the organization of cytoplasmic actin bundles but also associates with the lumenal plasma membrane of endothelial cells^21,22^. This state of knowledge indicates that paralemmins can functionally interact with both, the plasma membrane and the actin cytoskeleton, possibly linking them. However, the molecular details and mechanisms remained unknown.

In the present study, we investigated interaction partners and Palm1 function in neurons. Combining yeast-2-hybrid (Y2H) assays and advanced fluorescence nanoscopy, we show that Palm1 is a component of the MPS that binds to the N-terminal region of βII-spectrin, next to its actin-binding site. Overexpression, knock-out, and rescue experiments demonstrate that Palm1 expression levels are sufficient to control the nanoscale organization of the MPS. We propose that Palm1 achieves this by modulating interactions between MPS proteins around the N-terminal region of βII-spectrin, and tightening the attachment of this multidomain junction to the plasma membrane.

## Results

### Palm1 binds βII-spectrin and is a component of the MPS at the actin-adducin rings

To gain insight into the functions of Palm1, we searched for interaction partners. Y2H screening of a mouse brain cDNA library identified 258 interacting prey clones, all encoding N-terminal regions of βII-spectrin between the CH2 and the fourth spectrin-homology repeat (Fig. 1A). This large number of clones constituted a closely staggered “deletion series”, allowing high-resolution mapping of the βII-spectrin sequence interval necessary for Palm1 binding. Particularly at the 5’-ends, prey clones often differ only by one codon in length. An interval of 47 amino acids (aa) was delineated as the sequence of minimal overlap (SMO) of Palm1-interacting clones: PDEK…IEKY (aa 261-307). This sequence, close to the actin-binding domain, comprises the C-terminal α-helix of the calponin-homology domain 2 (CH2) and its linker with the first spectrin repeat (SR1).

**Figure 1:**
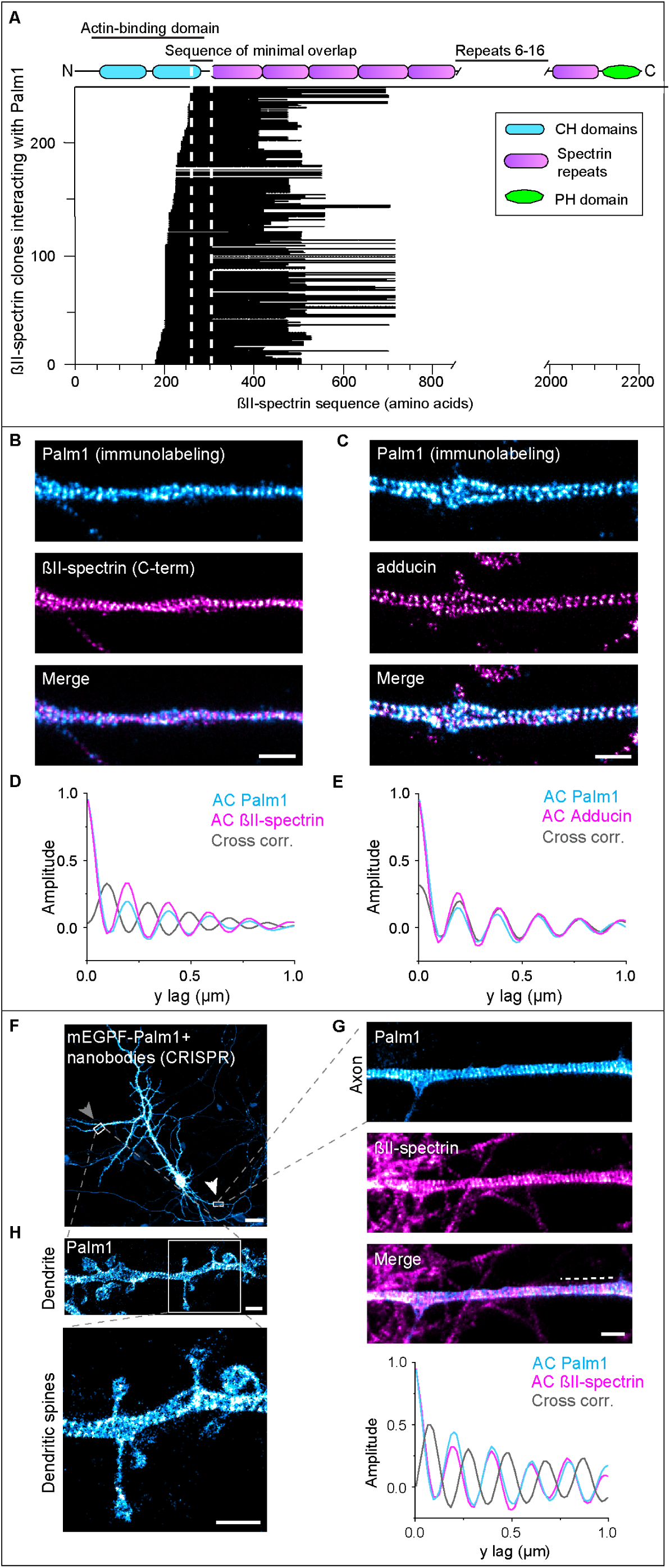
Palm1 is a MPS component. (A) Palm1-interacting prey clones aligned with the βII-spectrin sequence. Each horizontal line represents one of 258 prey clones. Vertical dashed lines indicate the SMO. CH: Calponin Homology, PH: Pleckstrin Homology. (B, C) Representative 2-color STED images of rat HPN (DIV 19, methanol fixation) immunolabeled for Palm1 and the βII-spectrin C-terminus (B) or adducin (C). Scale bars: 1 µm. (D) Average AC and CC analyses of Palm1 and βII-spectrin of n=27 axons from N=3 independent neuronal cultures. (E) As D for Palm1 and adducin (n=20, N=3). (F) Confocal image of a rat HPN (DIV 19, PFA fixation) expressing endogenous Palm1 tagged with mEGFP and detected by using a nanobody against mEGFP. Arrows point at regions displayed in G and H. Scale bar: 25 µm. (G) Close-up STED image of the axon co-immunostained for βII-spectrin. Scale bar: 1 µm. Dashed line on the merged image indicates the region on which the AC and CC analyses shown at the bottom have been performed. (H) Close-up STED images of a dendrite with dendritic spines. Scale bars: 1 µm.

The interaction with βII-spectrin, a key component of the neuronal MPS^2^ suggested that Palm1 might also be part of the MPS. Therefore, we investigated the nanoscale organization of Palm1 in mature rat hippocampal primary neurons (HPN) by indirect immunofluorescence (IF). Two-color STED nanoscopy revealed a periodic arrangement of Palm1 along the axons, alternating with the C-terminus of βII-spectrin (from now on, briefly “βII-spectrin”) (Fig. 1B) and colocalizing with adducin (Fig. 1C). This was quantified using 2D autocorrelation (AC) and cross-correlation (CC) analyses (Fig. 1D, E). AC revealed that Palm1 peaks with a period of ∼190 nm, characteristic of MPS components, while CC confirmed the out-of-phase and in-phase periodicity with βII-spectrin and adducin, respectively.

To exclude possible artifacts of Palm1-IF, we confirmed these observations by visualizing endogenously tagged Palm1 (Fig. 1F). CRISPR/Cas9 was used to knock-in mEGFP at the N-terminus of Palm1 in HPN. Also with this approach, Palm1 showed a clear periodic pattern along the axon, alternating with βII-spectrin, as confirmed by CC analysis (Fig. 1G). mEGFP-Palm1 also exhibited stretches of periodic organization along dendritic shafts and was seen in the necks and heads of dendritic spines (Fig. 1H). Thus, both Y2H interaction and STED nanoscopy indicate that Palm1 is a novel component of the MPS, binding to the N-terminus of βII-spectrin near its actin-binding site, and co-localizing at STED resolution with the actin/adducin rings.

### Palm1 populates distal neurites before MPS assembly, but incorporates into the MPS at a later stage

Next, we explored Palm1 subcellular distribution and nanoscale organization during development in culture. HPN at days *in vitro* (DIV) 1, 3, 5, 12, and 19 were co-stained for Palm1, βII-spectrin, and the proximal axon (AIS) marker ankyrinG (ankG)^23^ (Fig. 2A). Confocal images detected Palm1 in all neuronal compartments already from DIV 1, including distal neurites which were still poorly populated by βII-spectrin (Fig. 2A, B). Palm1 remained ubiquitously and evenly expressed along the axon during neuronal development in the middle (up to 40 µm after the AIS) and distal axon (>40 µm after the AIS), whereas βII-spectrin showed a strong proximal-to-distal gradient up to DIV 12 (Fig. 2A, D, F). However, parallel to βII-spectrin^13^, during HPN maturation Palm1 levels showed a reduction in the AIS and were complementary to ankG, as seen by the fluorescence signal intensities, a proxy for local protein concentrations (Fig. 2A, C, D, F). Thus, the axonal distribution of Palm1 largely parallels that of βII-spectrin, but Palm1 populates distal regions earlier than βII-spectrin, before the MPS assembly.

**Figure 2:**
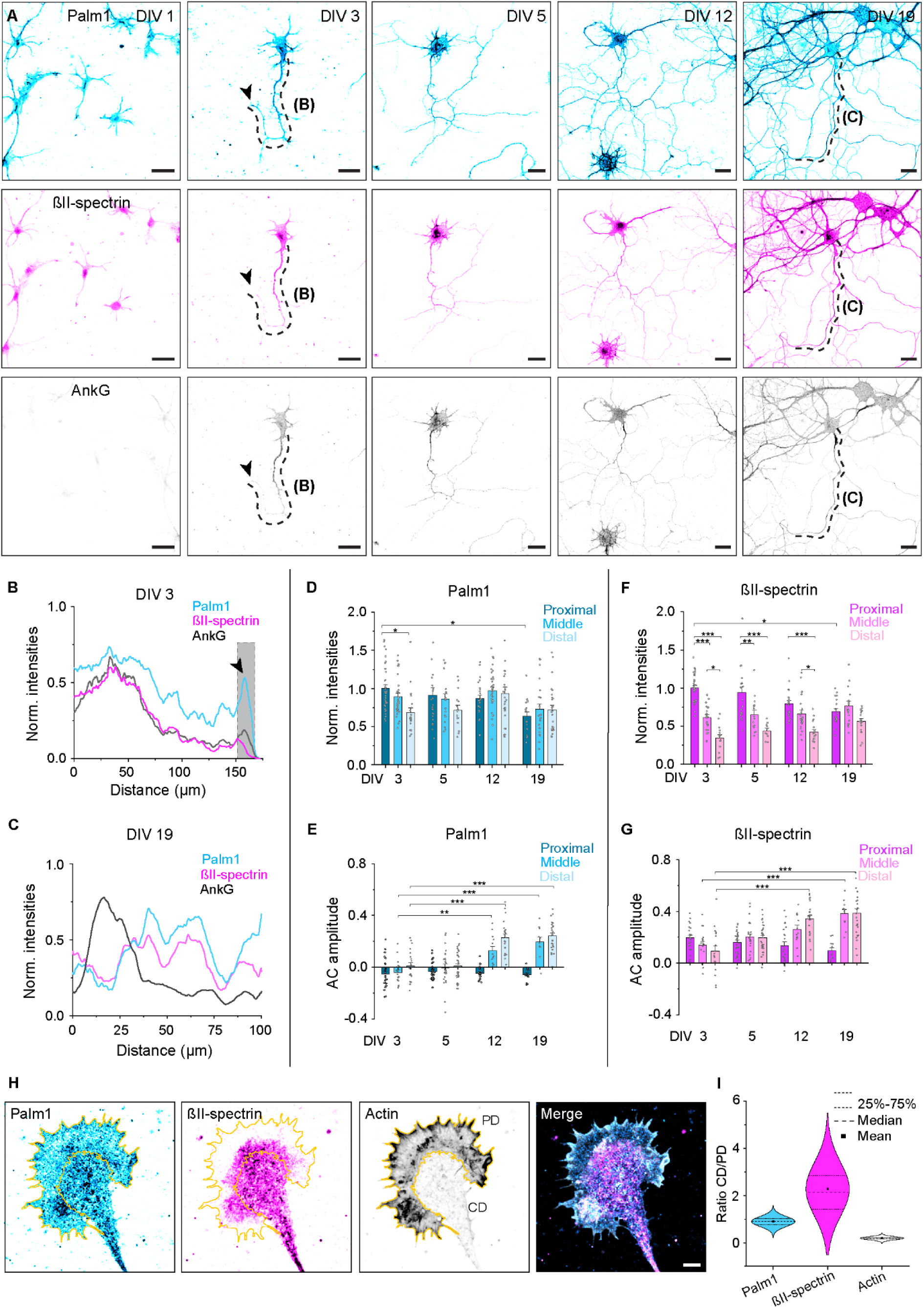
Palm1 populates distal axonal regions before βII-spectrin but is then incorporated into the MPS. (A) Representative confocal images of rat HPN (DIV 1, 3, 5, 12, and 19, methanol fixation) immunolabeled against Palm1, βII-spectrin, and ankG. Scale bars: 25 µm. Shown are the maximum intensities projection of 5 z-stacks. Black arrows point at the neurite end/growth cone. (B-C) Normalized (A.U., arbitrary units) and smoothed (50 values) fluorescence intensities of Palm1, βII-spectrin, and ankG along the axons indicated by the dashed lines on the representative image at DIV 3 (B) and at DIV 19 (C). Gray area in (B) highlights the typical enrichment of Palm1 at neurite end/growth cone. (D-G) Normalized fluorescence intensities (A.U.) and AC analyses of Palm1 (D, E) and βII-spectrin (F, G) along the proximal, middle, and distal axon at different DIVs. Axons analyzed for (D) and (E) in the proximal/middle/distal region: DIV 3: 35/31/21; DIV 5: 23/29/15; DIV 12: 25/33/26; DIV 19: 17/22/26. Axons analyzed for (F) and (G): DIV3: 24/18/20; DIV 5: 29/26/32; DIV 12: 22/13/23; DIV 19: 12/11/27. All from N=3. Statistical analyses: One-way ANOVA. p-values in file S2. Histograms show mean ± SEM. (H) Representative image of the growth cone of a HPN (DIV 2, PFA fixation), immunolabeled for Palm1 and βII-spectrin, and phalloidin-labeled for F-actin. Scale bar: 5 μm. (I) Fluorescence intensities ratio between the central (CD) and the peripheral (PD) domains of growth cones, segmented according to the phalloidin signal. Growth cones analyzed: n=56, from N=3.

To investigate whether also the nanoscale organization of Palm1 parallels that of βII-spectrin during axonal development, we analyzed the difference in the AC amplitude between the first peak at 190 nm and the average of the first 2 valleys (at 95 nm and 285 nm)^13^. AC amplitude is a measure of the periodicity quality: the higher the value, the more periodic the MPS nanoarchitecture. The negative AC amplitude indicates that Palm1 featured only an occasional and poor periodicity in all axonal regions in young neurons, when βII-spectrin periodicity is already evident and its AC amplitude positive (Fig. 2E, G). Palm1 acquired a long-range periodicity only from DIV 12, and only in the middle and distal axons. In the AIS, no Palm1 periodicity was observed regardless of the developmental stage and unlike βII-spectrin (Fig. 2E, G). Therefore, the time course of Palm1 nanoscale organization lags behind βII-spectrin. Furthermore, Palm1 periodicity does not correlate with its local concentration since in immature neurons, where Palm1 levels are already high, the protein is not periodically organized (Fig. 2D, E). A shift in Palm1 nanoscale organization occurs around DIV 12, when it becomes incorporated into the already established MPS although its local concentrations are constant. Similar results of Palm1 distribution and periodicity were obtained in mouse HPN (Fig. S1A-D).

Of note, in growth cones, Palm1 was evenly distributed over both the central and peripheral domains, including filopodia, whereas βII-spectrin and actin were enriched in the central and peripheral domains, respectively (Fig. 2H, I). Thus, growth cones may display, in a nutshell, Palm1 in two roles: a more stable (βII-spectrin-associated) as well as a more dynamic and more peripheral (βII-spectrin-independent) cell-biological context.

### Both Palm1 splice variants are co-expressed in neurons, promote neurite branching, and display reduced intra-axonal mobility

Two splice variants of Palm1 (full-length Palm1 and Palm1 without exon 8, Palm1ΔEx8) were identified in brain homogenates^17^. As these splice variants might be differentially expressed in neurons vs. glia, we wondered: (1) Are full-length Palm1 and Palm1ΔEx8 co-expressed in neurons? (2) Is the developmental change in the nanoscale organization of axonal Palm1 linked to a shift in differential splicing? We therefore extracted mRNA at different DIVs from HPN cultured in the presence of growth inhibitors for glial cells, and analysed the expression levels of both splice variants by RT-qPCR (Fig. 3A). Both splice variants were indeed co-expressed in HPN at similar levels. Whereas the expression of the Palm1 mRNA was unchanged between DIV 3-20, the Palm1ΔEx8 mRNA might display a slight biphasic developmental trend.

**Figure 3:**
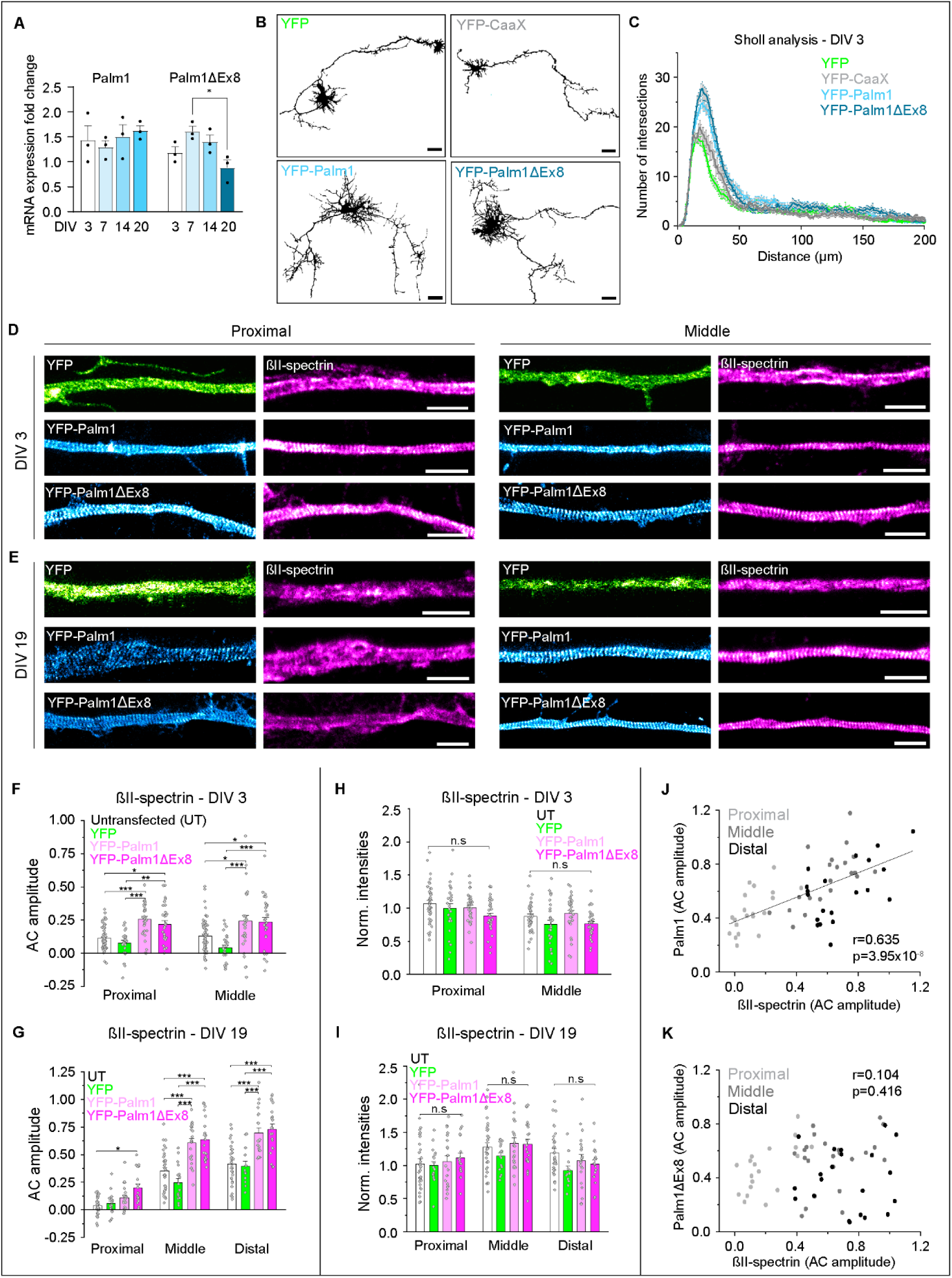
Overexpression of both Palm1 splice variants increases the complexity of neuronal morphology and enhances βII-spectrin periodicity. (A) mRNA expression levels of Palm1 and Palm1ΔEx8 during development of HPN. Fold change relative to the housekeeping genes. (B) Representative confocal images of rat HPN (DIV 3, PFA fixation) electroporated with plasmids encoding either YFP, YFP-CaaX, YFP-Palm1, or YFP-Palm1ΔEx8. Scale bars: 25 µm. (C) Sholl analysis of neurons overexpressing the indicated constructs. Intersections were counted every 1 µm. Cells analyzed: YFP: n=34, YFP-CaaX: n=24, YFP-Palm1: n=31, YFP-Palm1ΔEx8: n=27. All from N=3. (D) STED images of rat HPN at DIV 3 overexpressing YFP, YFP-Palm1 and YFP-Palm1ΔEx8, and endogenous βII-spectrin. Proximal (left) and middle (right) regions of the same axon are shown. YFP was detected using nanobodies. (E) Same as (D) but for DIV 19. Scale bars: 2 µm. Corresponding axons shown in Fig. S3. (F, G) AC amplitude analysis of endogenous βII-spectrin along different axonal regions in untransfected neurons (UT), or after overexpression of YFP, YFP-Palm1 and YFP-Palm1ΔEx8 at DIV 3 (F) and DIV 19 (G). (H, I) Normalized fluorescence intensities (A.U.) of βII-spectrin along the same axonal regions measured in (F) and (G), respectively. (J, K) Correlation scatter plots of the periodicity of βII-spectrin vs. Palm1 (J) or Palm1ΔEx8 (K). r, Pearson’s r coefficient; p, p-value. Axons analyzed in the proximal/middle region in (F) and (H): WT: 63/62; YFP: 27/27; YFP-Palm1: 31/30; YFP-Palm1ΔEx8: 31/31; and in (G) and (I-K): WT: 32/35/30; YFP: 18/17/13; YFP-Palm1: 20/21/20; YFP-Palm1ΔEx8: 19/23/21. All from N=3. Statistical analyses: One-way ANOVA; p-values in file S2. Histograms show mean ± SEM.

To test whether the Palm1 splice variants have specific roles in regulation of neuronal morphology or MPS incorporation rates, we overexpressed them. In immature neurons (DIV 3) they both induced an increased branching of neurites compared to controls (cytosolic YFP and membrane-anchored YFP-CaaX^24^), and Palm1ΔEx8 tended to have a more prominent effect (Fig. 3B-C). Soma morphology was unaffected (Fig. S2A-B). In mature neurons, both splice variants displayed a reduced motility in fluorescence recovery after photobleaching (FRAP) experiments, indicating impaired diffusion due to MPS incorporation (Fig. S2C-E), but no significant mobility differences. Therefore, both splice variants are expressed in neurons, but did not differ markedly in the functional parameters tested here.

### Palm1 overexpression enhances MPS periodicity without altering the local concentrations of MPS components

Next, we examined whether Palm1 overexpression impacts the MPS organization. Using STED nanoscopy, we analyzed the nanoarchitecture of both Palm1 and βII-spectrin along different axonal regions of neurons electroporated at DIV 0. In contrast to the non-periodic pattern of endogenous Palm1 in immature control cells, overexpressed YFP-Palm1 and YFP-Palm1ΔEx8 both displayed a long-range periodicity already at DIV 3 in the proximal and middle axon, confirmed by AC amplitude analysis (Fig. 3D, S2F, S3). Concomitantly, the periodicity of βII-spectrin was enhanced compared to untransfected cells or neurons transfected with YFP (Fig. 3F, and S3A). Also YFP-Palm1 and YFP-Palm1ΔEx8 overexpression in mature neurons (DIV 19), transfected at DIV 5, yielded high periodicities of YFP-Palm1 and YFP-Palm1ΔEx8 and significantly increased the periodicity of endogenous βII-spectrin, along proximal but especially middle and distal axons, and also along dendritic shafts (Fig. 3E, 3G, S2G, and S3B-D). The AC amplitude of YFP-Palm1 was slightly higher in the distal axons than YFP-Palm1ΔEx8 (Fig. S2G). In the AIS, albeit increased, Palm1 and βII-spectrin periodicities were lower compared to the other axonal regions (Fig. 3G, S2G, and S3B). In all conditions and cellular regions, recombinant Palm1 (like endogenous Palm1) alternated with βII-spectrin (Fig. S3A-D). We conclude that both recombinant Palm1 splice variants integrate into the MPS, and their increased levels enhance βII-spectrin periodicity in immature and mature neurons; in proximal, middle and distal axons; and in dendrites.

Previous studies showed that the local concentration of βII-spectrin promotes the formation of the MPS^13^. Since the overexpression of Palm1 enhanced the periodicity of βII-spectrin, we investigated whether this is due to an increased recruitment of βII-spectrin. We measured the fluorescent signal along the same axonal regions where the periodic rearrangement of βII-spectrin was analyzed. However, the local levels of βII-spectrin were unaffected by Palm1, independent of the neuronal developmental stage and axonal region (Fig. 3H-I). Hence, Palm1 overexpression did not lead to an increase in βII-spectrin protein levels, and no correlation between the local intensities of βII-spectrin and its periodicity was detected (Fig. S3E-F).

Comparing the periodicities (AC amplitudes) of YFP-Palm1 and βII-spectrin, a positive linear correlation was found: if in an individual axon Palm1 periodicity is low, also βII-spectrin periodicity is, whereas axons with highly periodic stretches of Palm1 correlate with highly organized βII-spectrin (Fig. 3J). For YFP-Palm1ΔEx8 no significant correlation was detectable at the single-axon level, although on average it also reorganized βII-spectrin (Fig. 3K). This difference might highlight different mechanisms of action of the two splice variants. Lastly, we tested whether the relationship between Palm1 and βII-spectrin levels and periodicity is mono- or bi-directional by transfecting neurons with βII-spectrin and immunolabeling for endogenous Palm1. However, no enhancement of Palm1 periodicity was observed upon βII-spectrin overexpression (Fig. S4A-B). Thus, Palm1 expression levels monodirectionally enhance βII-spectrin periodicity without altering its local concentrations. Similar to βII-spectrin, also adducin and ankB showed more robust periodic patterns along the middle and distal axons upon overexpression of YFP-Palm1, though not in the AIS, and unaltered local concentrations (Fig. S4C-J). Together, these data demonstrate that the overexpression of both YFP-Palm1 and YFP-Palm1ΔEx8 regulates the MPS along the middle and distal axons by enhancing the degree of its periodicity. Crucially, this effect is not due to an increase in the local concentrations of the MPS components βII-spectrin, adducin, or ankB, and it is monodirectional, with Palm1 driving βII-spectrin periodicity but not vice-versa.

### Depletion of Palm1 reduces the periodicity of the MPS and modifies electrophysiological parameters

Since the overexpression of Palm1 enhanced the periodic organization of the MPS, we expected that the depletion of the protein has the opposite effect. Hence, we analyzed HPN from constitutive Palm1-KO mice (Fig. S5A). We initially analyzed the development of Palm1-KO neurons from DIV 1 until DIV 19 by confocal microscopy. Palm1-KO cells exhibited a delayed development at DIV 1, with twice as many cells as the WT counterparts at stage 1 (soma surrounded by filopodia)^25^. At DIV 3, almost all neurons reached stage 3 (one neurite already outgrows the others) although the axonal specification was still lagging behind (lower βII-spectrin and ankG intensities) (Fig. S5B-D). Somata of Palm1-KO neurons were enlarged throughout development (Fig. S5E), and Sholl analysis at DIV 3 showed a slightly lower arborization degree compared to WT neurons (Fig. S5F).

We then investigated the nanoscale organization of the MPS along the axons of Palm1-KO mouse hippocampal neurons. Compared to WT neurons, βII-spectrin periodicity along middle and distal axons of Palm1-KO neurons was strikingly reduced in mature HPN (Fig. 4A-D). This effect was visible already at DIV 3 (Fig. S5G), *i.e*. even before the onset of Palm1 periodicity in WT neurons (Fig. 2E), and it may be cause or consequence of the developmental delay of immature Palm1-KO neurons. Importantly, βII-spectrin abundance in mature neurons remained unaltered, as shown by the local fluorescence intensities and Western blot analysis (Fig. 4E and S6A-B). Hence, the differences in MPS organization in mature Palm1-KO neurons cannot be ascribed to reduced βII-spectrin levels. Also the periodicities of adducin and ankB were reduced in the middle and distal axons of mature Palm1-KO neurons, whereas their local concentrations were unaffected (Fig. S6C-L). These observations indicate that, in the absence of Palm1, the MPS components βII-spectrin, adducin and ankB populate axons normally, but their periodic arrangement is severely compromised especially in middle and distal axons. As changes in the MPS organization can influence neuronal electrophysiology^26^ we explored electrophysiological parameters of mature Palm1-KO neurons. These cells exhibited a reduced miniature excitatory postsynaptic current (mEPSC) frequency (Fig. S7A-B). STED imaging of PSD-95 (a scaffold protein and marker of the postsynaptic density)^27^ revealed fewer but larger and brighter PSD-95 puncta along the dendritic shafts (Fig. S7C-E), suggesting that the observed decrease in mEPSC frequency is due to an altered postsynaptic compartment, in agreement with previous observations^19,28^. Palm1-KO neurons also exhibited a reduced rheobase (Fig. S7F-G), indicating higher excitation probability (*i.e.* a lower current suffices to trigger an action potential at the AIS). Higher excitability was also indicated by higher firing rates in Palm1-KO neurons upon somatic current injections (Fig. S7H-I). Changes in these parameters may reflect altered membrane excitability because of changes in MPS-related aspects, such as AIS position, structure, and membrane compactness, or reduced neurite branching. Together, these results show that Palm1 depletion not only impacts the organization of the MPS, but also compromises the intrinsic physiology of neurons and their ability to regulate postsynaptic compartments.

**Figure 4:**
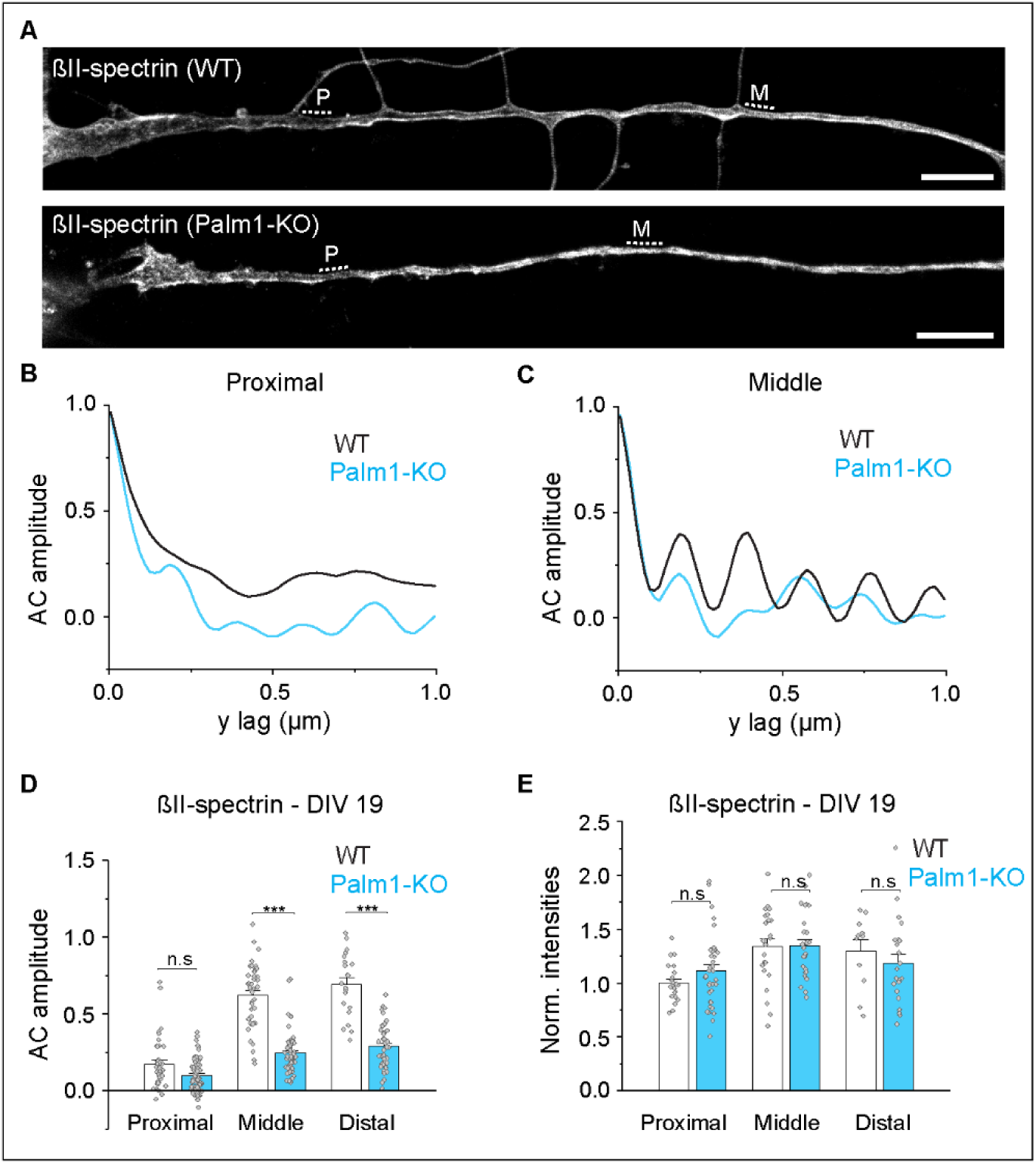
Palm1-KO mature neurons have a disorganized MPS. (A) Representative STED images of βII-spectrin nanoscale organization along the proximal and middle axon in mouse WT and Palm1-KO neurons (DIV 19). Scale bars: 5 µm. (B-C) AC amplitudes calculated from the regions indicated by the dashed lines in (A) (P: proximal; M: middle). (D) AC amplitudes analysis and (E) normalized fluorescence intensities (A.U.) of endogenous βII-spectrin of the same regions in the proximal and middle axons of WT and Palm1-KO neurons. Axons analyzed in the proximal/middle/distal region in (D): WT: 38/43/21; Palm1-KO: 70/58/42; and in (E): WT: 22/23/11; Palm1-KO: 36/27/22. All from N=3. Statistical analyses: One-way ANOVA; p-values in file S2. Histograms show mean ± SEM.

### Palm1 re-introduction into KO neurons rescues and enhances MPS periodicity

The constitutive Palm1-KO dramatically reduced the MPS periodicity. To clarify whether the short-term re-introduction of Palm1 is sufficient to restore it, we performed rescue experiments in which YFP-Palm1 or YFP-Palm1ΔEx8 were transiently overexpressed in Palm1-KO neurons. STED imaging showed that only neurons transfected with either Palm1 variant exhibited a clear and long-range periodic βII-spectrin structure in virtually all neurites (Fig. 5A-B). Again, the local concentrations of βII-spectrin were unaffected (Fig. 5C). Linear correlations with the periodicity of βII-spectrin were observed for both splice variants, more pronounced with YFP-Palm1 than with YFP-Palm1ΔEx8 (Fig. 5D-E). These correlations were more marked in this experiment than in WT neurons (Fig. 3J-K), where the background of endogenous Palm1 might dilute the effect. These rescue results confirm that Palm1 is a powerful regulator of the MPS nanoscale architecture.

**Figure 5:**
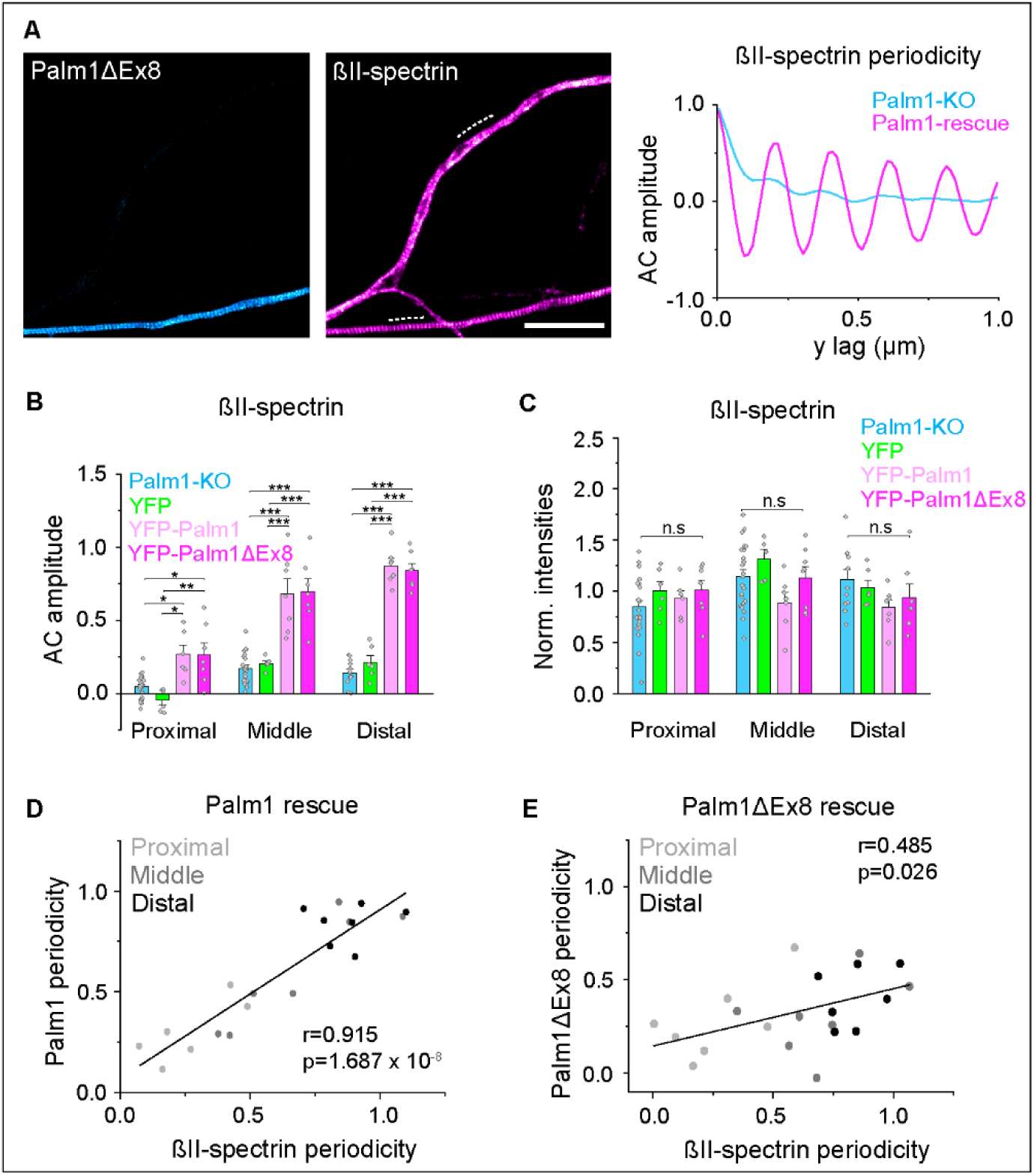
Palm1 re-introduction into Palm1-KO neurons rescues and enhances the periodic organization of the MPS. (A) Representative STED images of recombinant YFP-Palm1 and endogenous βII-spectrin fluorescence periodicity displaying an untransfected axon lacking Palm1 (top neurite) and an axon rescued by overexpression of YFP-Palm1ΔEx8 (bottom neurite; DIV 13, PFA fixation). AC analysis along the dashed lines shows the enhanced periodic pattern of βII-spectrin after Palm1ΔEx8 overexpression. Scale bar: 4 µm. (B) AC amplitudes analysis of βII-spectrin along different axonal regions in untransfected Palm1-KO neurons, or after overexpression of YFP, YFP-Palm1 or YFP-Palm1ΔEx8 (DIV 13, transfection at DIV 5). (C) Normalized intensities (A.U.) of endogenous βII-spectrin along the same axonal regions analyzed in (B). (D) Correlation scatter plot of βII-spectrin periodicity vs. Palm1 or (E) Palm1ΔEx8. r, Pearson’s r coefficient; p, p-value. Axons analyzed for B-E in the proximal/middle/distal region: Palm1-KO: 23/23/12; YFP: 6/6/6; YFP-Palm1: 6/7/7; YFP-Palm1ΔEx8: 7/7/7. All from N=1. Statistical analyses: One-way ANOVA; p-values in file S2. Histograms show mean ± SEM.

### The “paralemmin sequence motif” is required for the MPS-binding and -remodelling activities of Palm1

The “paralemmin motif” is a sequence feature of 11 aa, conserved throughout evolution in all paralemmin isoforms and vertebrate species^18^ (in Palm1: K46…L56), and must therefore be important for paralemmin function. Introducing the single missense mutation W54A into Palm1 abolished the integration of overexpressed YFP-Palm1(W54A) into the MPS of WT neurons, leaving the mutant Palm1 localized at the plasma membrane but lacking periodicity (Fig. 6A, B). Furthermore, the enhancement of βII-spectrin-periodicity caused by overexpression of non-mutant YFP-Palm1 failed to occur with YFP-Palm1(W54A) (Fig. 6C). The baseline periodicity of βII-spectrin remained unaffected, suggesting that the disabled YFP-Palm1(W54A) protein cannot associate with and remodel the MPS. These findings demonstrate that the “paralemmin motif”, particularly its tryptophan residue W54, is essential for Palm1 recruitment to the MPS and the enhancement of MPS periodicity.

**Figure 6:**
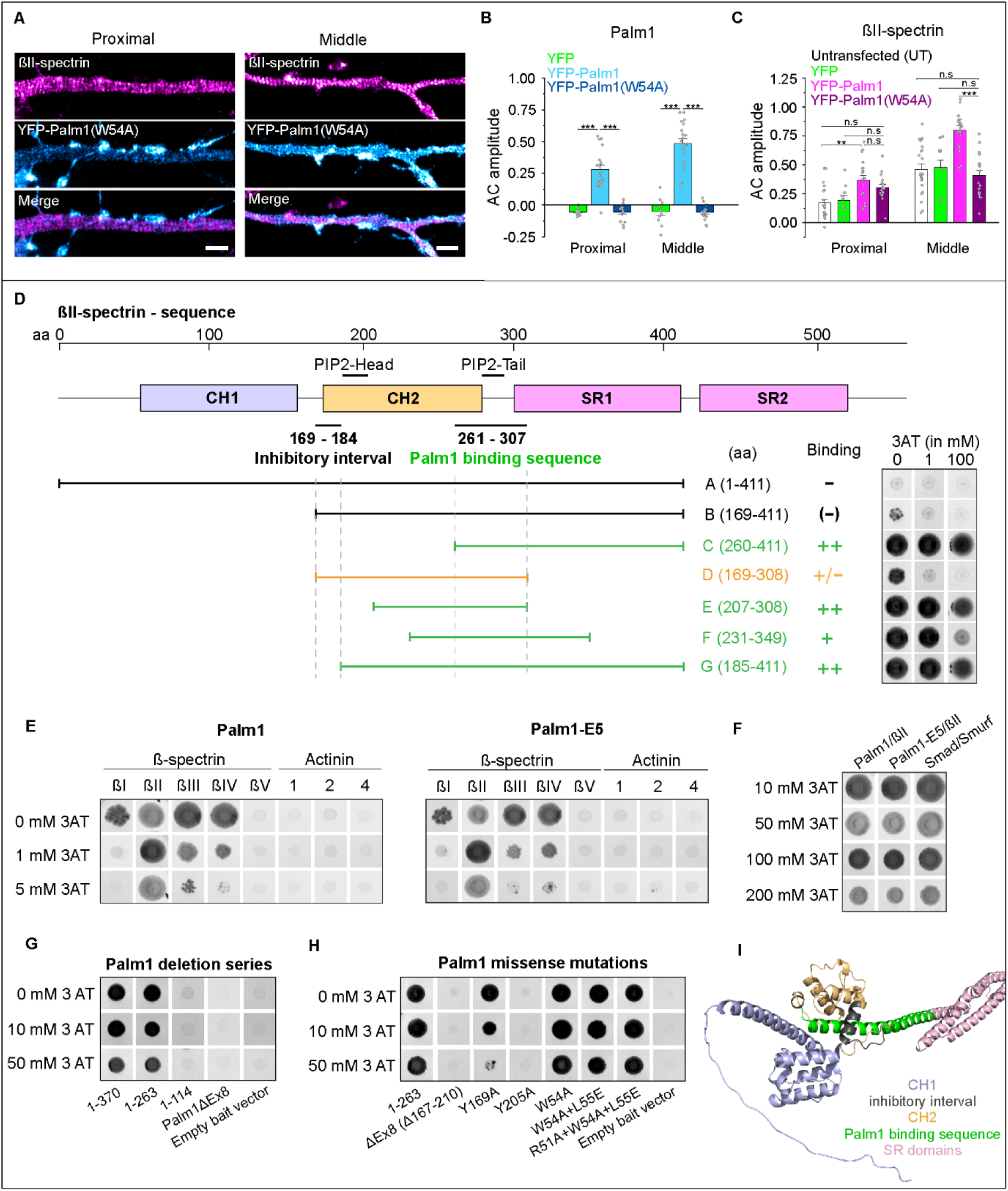
Molecular features involved in the interactions between the MPS, Palm1, and β-spectrin. (A) The W54A mutation in Palm1 abolishes MPS integration and remodeling. STED-IF images of rat HPN at DIV 14 (PFA fixation) overexpressing YFP-Palm1(W54A), and endogenous βII spectrin along the proximal (left) and middle axon (right). Scale bar: 1 µm. (B) AC amplitude analyses of YFP, YFP-Palm1, YFP-Palm1(W54A) and (C) βII-spectrin along different axonal regions in untransfected neurons (UT); or after electroporation of YFP, YFP-Palm1 or YFP-Palm1(W54A) (DIV 13-15). Axons analyzed for B-C in the proximal/middle region: YFP: 10/11; YFP-Palm1: 19/20; YFP-Palm1(W54A): 16/18; Untransfected: 20/21. All from N=2-3. Statistical analyses: One-way ANOVA; p-values in file S2. Histograms show mean ± SEM. (D) Schematic of the N-terminal region of βII-spectrin, depicting the CH domains and the first two SR domains. The sequence necessary for Palm1 binding (aa 261-307) and the inhibitory interval (aa 169-184) blocking Palm1/βII-spectrin interaction are indicated, as well as the collinear PIP2-binding sites in actinin-2. Below, the βII-spectrin prey constructs A-G are schematically displayed together with the respective yeast spot colonies detecting these interactions (right). (E) Comparative testing of Palm1 and the phosphomimetic-mutant Palm1-E5 for interaction selectivity with five β-spectrin and three actinin isoforms as preys. (F) Interaction of Palm1 and Palm1-E5 with βII-spectrin is unaffected by high 3AT concentrations. Smad/Smurf: positive control. (G) Yeast spot colonies of a C-terminal Palm1 deletion series and Palm1ΔEx8 as baits. (H) Yeast spot colonies of truncated Palm1(1-263), its ΔEx8 splice variant, and several missense mutants as baits. Yeast colonies images are shown in the negative for better visualization. In experiments D, G and H, a complete 3AT concentration series (0, 1, 2, 5, 10, 20, 50, 100, 200 mM) was tested, but for clarity only selected concentrations with the most informative growth patterns are shown. In experiments G-H, βII-spectrin construct G was used as prey. (I) Rendering of βII-spectrin N-terminal structure highlighting the relevant domains and sequences regulating Palm1 binding (Video S1). Structure was predicted by AlphaFold2 (Q62261, aa 1-529).

### Molecular features of β-spectrin involved in Palm1 binding

To explore the molecular mechanisms through which Palm1 exerts its function at the MPS, we returned to the Y2H system. The high-resolution deletion series constituted by the Palm1-interacting βII-spectrin clones of our Y2H screen had narrowed down their sequence of minimal overlap (SMO) to 47 aa (aa 261-307) (Fig. 1A). To dissect the involvement of βII-spectrin structural elements in Palm1 binding in further detail, we constructed and tested a set of βII-spectrin preys (Fig. 6D). For a semi-quantitative measure of interaction strength, yeast were grown on a concentration series of 3-amino-1,2,4-triazol (3AT), a competitive inhibitor of the histidine biosynthesis pathway. A striking feature of the deletion series of Fig. 1A was that no prey clones reaching further N-terminal than aa 185 were picked up, suggesting that the N-terminal ∼184 aa of βII-spectrin block its interaction with Palm1 in the Y2H assay. This was confirmed in the present experiment by the negative outcome with construct A (aa 1-411, including both CH domains) and weak residual activity of construct B (aa 169-411, retaining the complete CH2 domain). Strikingly, the deletion of a mere further 16 N-terminal aa in construct G (aa 185-411) restored binding activity. βII-spectrin constructs C, E and G (all lacking the inhibitory interval) displayed indiscriminable interaction strengths, whereas that of construct F (aa 231-349) was slightly weakened. Construct D (aa 169-308) retained partial interaction activity in spite of reaching up to aa 169; as it lacks most of the SR1 domain at its other end, this suggests that the SR1 domain participates in the inhibition of Palm1 binding by aa 169-184 (which corresponds to the first α-helix of the CH2-domain). Thus, we have mapped the Palm1-binding site on βII-spectrin with high precision, placing it in a location of potential interplay with several other molecules at the junction of βII-spectrin with actin, adducin and the calmodulin-like domains of αII-spectrin^5,29–31^.

β-Spectrin isoforms βI-βV and actinin isoforms 1-4 share homologous domain architectures, with N-terminal CH1-CH2 actin-binding domains followed by SR repeats^29^. We therefore explored, (1) whether Palm1 binds only βII-spectrin, or might cross-react with these related proteins; and (2) as Palm1 is multiply phosphorylated in neurons, whether phosphomimetic mutations of known phosphorylation sites of Palm1 influence the binding to β-spectrin/actinin isoforms. New prey constructs were modeled on βII-spectrin construct G (aa 185-411), which encompasses the CH2 domain (except its inhibitory first α-helix), the SR1 domain, and the linker between them. The high sequence similarity between the β-spectrins and actinins allowed a very accurate alignment and the design of preys of the same length (227-228 aa, see Methods). These preys were matched with Palm1, and with the phosphomimetic mutant Palm1E5 (carrying mutations of ten serine or threonine residues, known to be phosphorylated in mouse brain, to glutamate). Palm1 interaction with the βII-spectrin prey enabled robust yeast growth unaffected by up to 100 mM 3AT, as robust as that of the Smad/Smurf positive control (Fig. 6E-F). Also βI-, βIII- and βIV-spectrin preys enabled good growth at 0 mM 3AT which was increasingly suppressed by 1-5 mM and abolished by 10 mM 3AT, whereas βV-spectrin and the actinin isoforms did not interact at all with Palm1 (Fig. 6E). These interactions were unaffected by the ten phosphomimetic mutations of Palm1E5. The specificity spectrum of Palm1 for β-spectrin and actinin isoforms correlates well with their sequence similarities: β-spectrin isoforms I, III and IV have 81-87% aa identity to the βII-spectrin SMO, whereas βV-spectrin and actinins 1-4 share only 40-45% aa identity with the βII-spectrin SMO sequence. Other, more distantly related proteins with CH1-CH2 tandem domains (dystrophin, utrophin, nesprin, MACF1, BPAG1, plectin, smoothelin, filamin) have even lower sequence similarities with the βII-spectrin SMO (13-26% identity, limited to the last CH2 α-helix) and were not tested here.

We conclude that βII-spectrin is the main target of Palm1, but Palm1 can also interact with βI-, βIII- and βIV-spectrin, albeit less avidly. The Palm1 – βII-spectrin Y2H interaction requires the last α-helix of the βII-spectrin CH2 domain as well as the adjacent CH2-SR1 linker sequence, and is unaffected by the ten phosphomimetic mutations tested here. Importantly, the CH1 domain and particularly the first α-helix of the βII-spectrin CH2 domain inhibit this interaction.

### Palm1 binds β-spectrin via the “core domain” encoded by differentially spliced exon 8

Having defined molecular features of β-spectrin involved in Palm1 interaction, we went on to identify the parts of Palm1 needed for binding βII-spectrin. A C-terminal deletion series (Fig. 6G) demonstrated that binding to βII-spectrin construct G (185-411) depends on the central Palm1 region (aa 115-263). This interval includes the “paralemmin core domain”, most of it encoded by the differentially spliced exon 8 (aa 167-210), and indeed Palm1ΔEx8 was also unable to interact with βII-spectrin. In the Y2H assay, deletions of longer sequence intervals may unspecifically affect interactions by perturbing the sterical arrangement of the hybrid protein complex, and we therefore probed the Palm1 sequence elements necessary for βII-spectrin interaction more specifically by introducing missense mutations. In the C-terminally truncated bait sequence Palm1(1-263) which retains full βII-spectrin interaction, we analysed several structural variants for their effect on the interaction with βII-spectrin construct G (Fig. 6H): deletion of exon 8; two individual missense mutations of tyrosine residues in exon 8 (Y169A or Y205A; conserved between Palm1, Palm2 and Palmd) and three cumulative missense mutations in the “paralemmin sequence motif” which are conserved between all four paralemmin isoforms (W54A; W54A+L55E; R51A+W54A+L55E). The deletion of exon 8 and the Y205A mutation completely abolished interaction even in the absence of 3AT. The Y169A mutation reduced interaction, progressively weakened from 2 mM 3AT and abolished by 50 mM 3AT. In contrast, the baits with mutations in the paralemmin motif around W54 displayed activities indiscriminable from the unmutated Palm1(1-263) bait and partially reduced only by 100-200 mM 3AT (not shown). We conclude that the paralemmin core domain largely encoded by exon 8, and not the “paralemmin sequence motif”, mediates the interaction between Palm1 and the βII-spectrin SMO, and that tyrosine residue Y205 is critical for this activity.

### Palm1 closely flanks adducin at the actin rings

According to the Y2H data, Palm1 binds close to the spectrin/actin junction, with which also adducin interacts^5,32^. STED nanoscopy had shown that Palm1 and adducin indeed co-localize, at a resolution of 30-60 nm (Fig. 1C). To determine the relative positions of Palm1 and adducin even more precisely, we applied MINFLUX, a light microscopy technique that allows the localization of molecules with single-digit nanometer precision^33^. Because the overexpression of YFP-Palm1 enhances the periodicity of the MPS without altering the concentrations of its components, we used this preparation as our model for MINFLUX imaging with DNA-PAINT^34^. 3D MINFLUX imaging clearly visualized the periodic organization of Palm1 (Fig. 7A, Video S1). A YZ projection of an axon segment along its main axis reveals a hollow structure, in accordance with Palm1 being membrane-associated and not cytosolic (Fig. 7B). To extract quantitative information from the 3D data, we unwrapped the axons and displayed the 3D information in 2D (Fig. 7C)^35^. This rendering revealed that Palm1 is arranged in relatively broad bands. The molecules were spaced between 6 nm for the 1^st^ Nearest Neighbor (NN, this value might include multiple localization of the same molecule), to 10.3 nm for the 2^nd^ NN, up to 16.5 nm for the 4^th^ NN (Fig. 7D). We then used exchange DNA-PAINT to simultaneously detected Palm1 and adducin. When unwrapping the dual-channel data of axons, we observed that adducin molecules were located in close proximity to Palm1, and in many cases, they were sandwiched by Palm1 (Fig. 7E, Video S1). The NN distance between adducin and Palm1 peaked between ∼5 and 20 nm, including the size of the labels of ∼5-10 nm^36^ (Fig. 7F). This distance is comparable to the width of F-actin filaments which constitute the rings (∼18 nm for the actin braids and ∼10 nm for the single filaments)^4^. In this way, MINFLUX positions Palm1 (specifically, its YFP-tagged N-terminus) at both sides of the actin-spectrin-adducin junctions with a quantification of physical proximity *in situ*.

**Figure 7:**
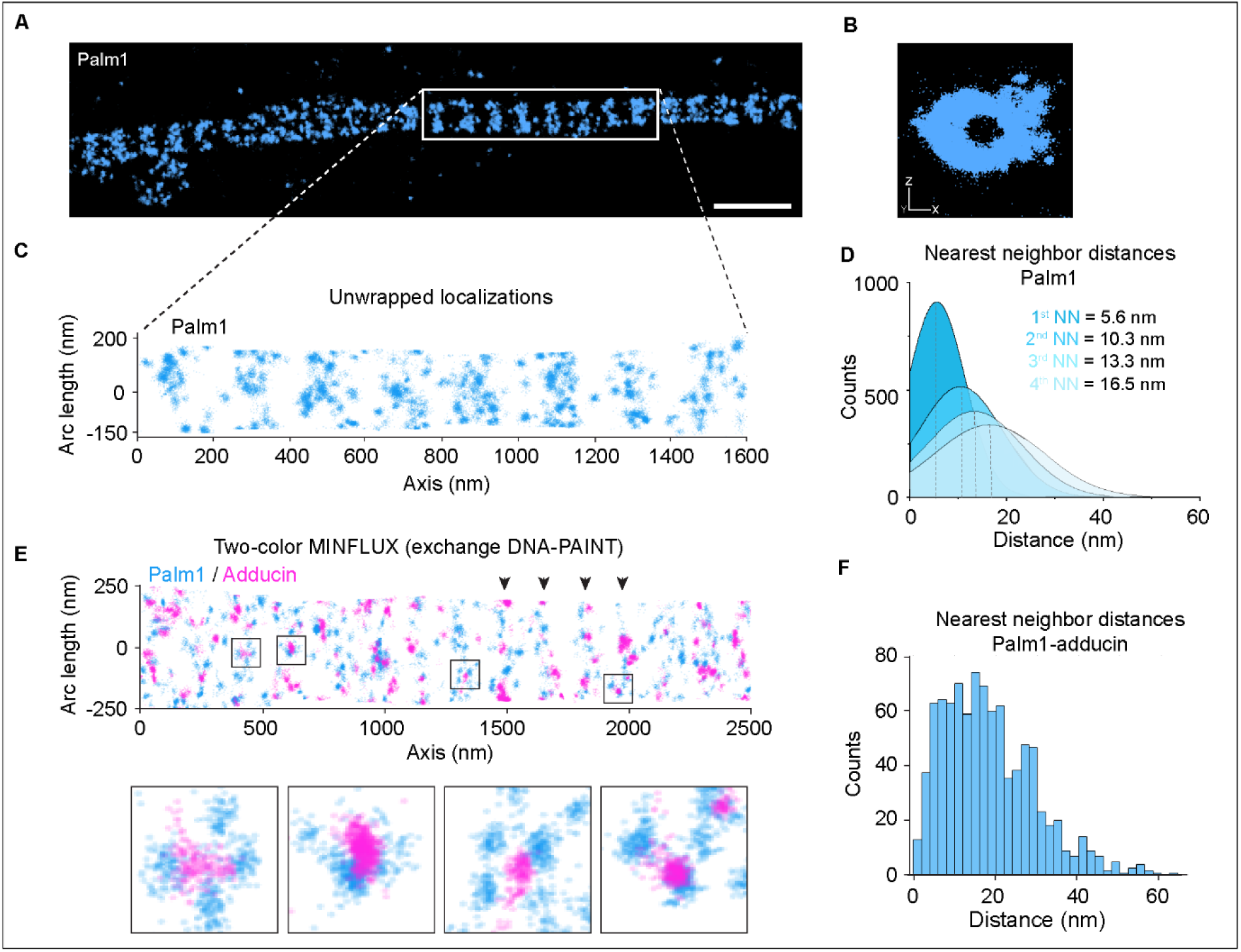
Palm1 proximity to adducin at the MPS. (A) Single-color 3D MINFLUX of periodic YFP-Palm1 along the axon of a mature neuron (DIV 18, PFA fixation). Scale bar: 500 nm. Palm1 was detected by using a nanobody against YFP in combination with DNA-PAINT. (B) YZ projection of the region highlighted in (A) shows a hollow structure, indicating that Palm1 localizes to the plasma membrane. (C) 2D projection of region indicated in (A). (D) First to fourth nearest neighbor (NN) analysis of Palm1 molecules. Data from 6075 trace ID (TID, *i.e.* individual localization bursts) from four MINFLUX images. (E) Two-color 3D MINFLUX (exchange DNA-PAINT) reveals clear periodic patterns (highlighted by black arrowheads) for both YFP-Palm1 (blue) and adducin (magenta). Black boxes indicate Palm1 doublets flanking adducin molecules displayed in the close-ups below. (F) NN distances of Palm1 and adducin obtained from 918 TIDs corresponding to the image shown in (E).

## Discussion

In this study, we identify Palm1 as a new component and regulator of the MPS. Palm1 binds to βII-spectrin at an N-terminal site comprising the end of the CH2 domain and the CH2-SR1 linker, and is concordantly observed *in situ* to closely flank adducin at the actin rings of the MPS. The periodicity of βII-spectrin, adducin and ankB, major structural components of the MPS, is drastically reduced and barely detectable in Palm1-KO neurons, whereas conversely, overexpression of recombinant YFP-Palm1 enhances the periodicities of all tested MPS components above those at WT Palm1 levels. Thus, the expression levels of Palm1 tune the degree of MPS periodicity, while leaving the local concentrations of βII-spectrin, adducin and ankB unaffected. Hence, Palm1 is the first component of the MPS primarily dedicated to controlling its periodicity (Fig. 8).

**Figure 8:**
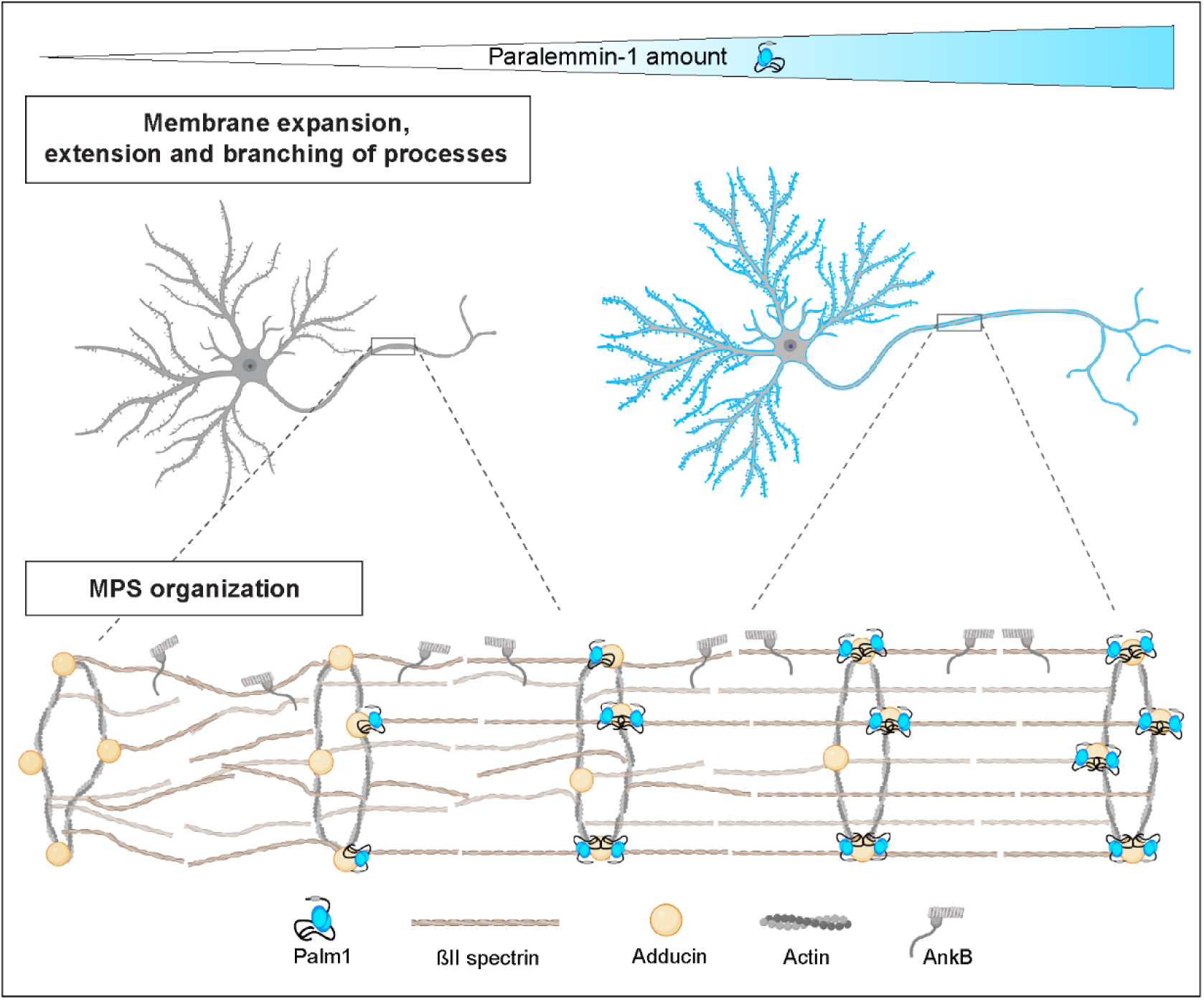
Visualization of Palm1-induced cellular and molecular manifestations in relation to its expression levels.

Three sequence regions, highly conserved between paralemmin isoforms in multiple animal species^18,38^, are hallmarks of this protein family. The N-terminal ∼120 aa are predicted to form an antiparallel coiled-coil in all four isoforms, and carry at their center a sequence signature termed the “paralemmin motif”, which in Palm1 centers around W54^18^. A second region of high homology encodes the “paralemmin core domain”, shared only by Palm1, Palm2 and Palmd. Alphafold predicts it to fold into a characteristic arrangement of a four-stranded β-sheet and a juxtaposed α-helix (https://alphafold.ebi.ac.uk/entry/Q9Z0P4); in Palm1, it can be abolished by differential splicing of exon 8 (https://alphafold.ebi.ac.uk/entry/Q542N8). The third homology region toward the C-terminus includes the “MIF (Met-Ile-Phe) motif”, again shared only by Palm1, Palm2 and Palmd, which is predicted by Alphafold to insert as a fifth strand into the β-sheet of the core domain. However, C-terminal truncation of Palm1(1-263) removes the third homology region without reducing the βII-spectrin interaction in Y2H.

Our present findings assign cell-biological or molecular functions to the first two homology regions. The “paralemmin motif” in the coiled-coil domain (particularly W54) is essential for Palm1 to be recruited to the MPS and enhance its periodicity. However, even up to three missense mutations in the paralemmin motif do not disrupt the Y2H interaction with βII-spectrin. The “core domain” (particularly Y205), abolished by differential splicing of Palm1 exon 8, is essential for the very avid interaction with the CH2-SR1 linker of βII-spectrin in the Y2H assay, but has little effect on the MPS-binding and remodeling activity of Palm1. How these two domains of Palm1 and their respective functional properties are mechanistically connected, however, is still unclear. The +/- exon 8 splice variants showed only gradual differences, or none at all, for several functional parameters after overexpression in HPN (neurite complexity, MPS periodicity, and intra-axonal mobility via FRAP). We propose that Palm1 interacts with several partner molecules around the actin – spectrin junction complex, perhaps in a sequence of mechanistic steps, orchestrating the assembly and regulation of this junction. The seminal findings of the present study pave the way for future research to comprehensively unravel the mechanisms through which Palm1 and its isoforms interact with the actin – spectrin cytoskeleton.

The promotion of MPS periodicity by Palm1 probably requires not only the “paralemmin motif”, but also the C-terminal membrane anchor, a cluster of three lipidated cysteine residues (”CaaX box”) which anchors it in detergent-resistant, lipid raft-like membrane microdomains^17,39^. This probably stabilizes the position of the Palm1-αII/βII-spectrin-actin-adducin junction, both vertically (in terms of plasma membrane distance) and horizontally (by impeding lateral diffusion within the plasma membrane). Deletion or mutation of the C-terminal CaaX motif abolishes the plasma membrane-remodeling activity of Palm1^17,19^, emphasizing the importance of the membrane anchor.

The position of the Palm1-binding site of βII-spectrin suggests a high potential for molecular interplay with the neighboring CH1 and CH2 domains, and additional binding partners of this region: the EF-hand domains of αII-spectrin, actin, adducin (which promotes the β-spectrin – actin interaction), and potentially protein 4.1 and the phosphoinositide PtdIns(4,5)P2 (PIP2). The first example for interplay, observed in our present experiments, is the identification of the first α-helix of the CH2 domain as an intramolecular inhibitor of Palm1 binding (Fig. 6D). Previously, removal of the corresponding α-helix in the erythrocyte βI-spectrin isoform enhanced the binding of protein 4.1R and additionally exposed a cryptic actin-binding activity of the CH2 domain^40^. As overexpressed YFP-Palm1 in neurons readily integrates into the MPS and reorganizes it (Fig. 3D-G), there must be mechanisms to relieve this block *in situ*. Several additional ligands converge on the C-terminal part of the βII-spectrin SMO sequence, *i.e.* the linker between the CH2 and SR1 domains. In βI-spectrin, this region binds the two C-terminal calmodulin-like domains of αI-spectrin^32^, amplifying the binding of the CH1-CH2 domains to F-actin,^30^ and an analogous inter-chain interaction occurs in actinin-2^41^. The phosphoinositide lipid PIP2 binds to, and functionally affects, both actinin-2 and βI-spectrin^40,42^. The binding site for the PIP2 head group has been located in the CH2 domain of actinin-2^42^ (a sequence conserved in βII-spectrin but outside the Palm1-binding site), and the PIP2 lipid tail was proposed to bind to the CH2-SR1 linker, overlapping with the Palm1-binding site (Fig. 6D)^43^.

In proteins containing tandem CH-domains, such as βII-spectrin, this actin-binding module appears to interconvert between a closed conformation, in which the CH1 and CH2 domains bind each other and exclude actin, and an open conformation which frees the CH1 domain for binding to actin^44,45^. Numerous heterozygous disease mutations have been identified in the CH2 domains of the human βII-spectrin, βIII-spectrin and actinin-4 genes^31,37,46–48^. Twenty-eight mutations in the human βII-spectrin gene were found in association with autosomal-dominant neurodevelopmental disorders^48^. Half of these mutations concentrate in the CH2 domain, where they seem to interfere with the CH1-CH2 closed conformation and thus de-regulate actin binding^48^. Five of these 28 βII-spectrin missense mutations lie within aa 268-275, *i.e.* within the Palm1-binding SMO, suggesting that these aa are crucial for the closed CH1-CH2 conformation, and that by binding to this region Palm1 might wedge into the CH1-CH2 interface, separate the two CH domains, and thus promote actin binding by CH1. In actinin, the first CH2 helix was shown to be part of an actin-binding interface, and the last CH2 helix to be involved in CH1-CH2 dimerization, consistent with potential interplay between actin binding to the former and Palm1 binding to the latter in βII-spectrin^49^.

Disease mutations associated with a neurological disorder (spinocerebellar ataxia) were also identified in the CH2-domain of βIII-spectrin, three of them in the sequence homologous to the Palm1-binding site of βII-spectrin^31^. For the neighboring L253P mutation of βIII-spectrin, enhancement of actin binding through opening of the CH1-CH2 conformation has been directly demonstrated and characterized as a plausible pathomechanism^31,46,47^. Finally, mutations in the human actinin-4 gene cause a severe renal disorder (focal segmental glomerulosclerosis, FSGS). Strikingly, eight out of 22 such mutations concentrate in the last α-helix of the CH2 domain, and seven of these affect aa residues which are identical in βII-spectrin and lie in βII-spectrin’s Palm1-binding SMO^24^. These genetic findings collectively emphasize the outstanding functional importance of the last α-helix of the CH2 domain of β-spectrin/actinin isoforms. As this molecular feature is part of the Palm1-binding site of βII-spectrin, we can expect that also Palm1 binding to it will have a marked functional effect. Future research will have to explore the complex molecular interplay at and around the βII-spectrin CH2 domain further, and how Palm1 is woven into it as an additional partner. Large parts of the Palm1 sequence are intrinsically unstructured (https://alphafold.ebi.ac.uk/entry/Q9Z0P4), and after recruitment of Palm1 to the junction complex, they may wrap around the neighboring proteins and stabilize the whole assembly, or they may recruit yet additional ligands to it.

Palm1 is highly phosphorylated, particularly in the brain, and its phosphorylation status is affected by synaptic activity^50^. Around 30 serine/threonine phosphorylation sites have been identified in mammalian Palm1 (*i.e.* nearly one in every 10 aa), mostly in the intrinsically unstructured sequences (www.phosphosite.org; N.G.M and M.W.K., unpublished). This suggests that Palm1 phosphorylation may mediate regulatory input into the MPS, e.g. during development or neuronal activity. The Palm1 – βII-spectrin Y2H interaction was not significantly affected by the ten phosphomimetic mutations which we tested (Fig. 6E-F). Therefore, the molecular and functional parameters affected by Palm1 phosphorylation still await identification.

Palm1 is expressed in many tissues and cell-types, though highest in brain where it is quite abundant (∼0.2% of total mouse brain protein; G. Hultqvist and M.W.K., unpublished). Therefore, the biological role of Palm1 must be broader, and its involvement in the MPS is only one aspect. In cells or subcellular locations where βII-spectrin is absent, it may instead interact with other binding partners (e.g. its other Y2H interactors βI-, βIII- or βIV-spectrin). The overexpression of Palm1 in cell-lines and neurons was previously observed to stimulate cell expansion and the extension of polymorphic cell processes and filopodia^17,19^. In our present experiments the overexpression of Palm1 promoted the number and branching of neurites, while the Palm1-KO delayed the early morphological differentiation of neurons. Palm1 and its isoforms may participate in many different aspects of the cell biology of neurons and other cell types also outside the MPS, e.g. in neuronal growth cones (Fig. 2H). The effects on several electrophysiological parameters and on postsynaptic compartments observed in Palm1-KO neurons also suggest this, although disentangling MPS-related and -independent components will be difficult. The MPS has been found in virtually every neuronal cell type from invertebrates (*i.e*. D. melanogaster or C. elegans) to vertebrates (chicken, mouse, rat, and humans)^1–3^ indicating that it is ancient and conserved during evolution. Paralemmins, however, exist only in vertebrates^18^ suggesting that they arose later in evolution as a means to modulate the membrane-associated skeleton in more complex organisms. In addition, considering the high degree of phosphorylation of Palm1 and its isoforms, it may contribute to the non-structural functions of the MPS, such as the mediation of cell signaling and cell-cell interactions.

## Materials and methods

**Table 1:**
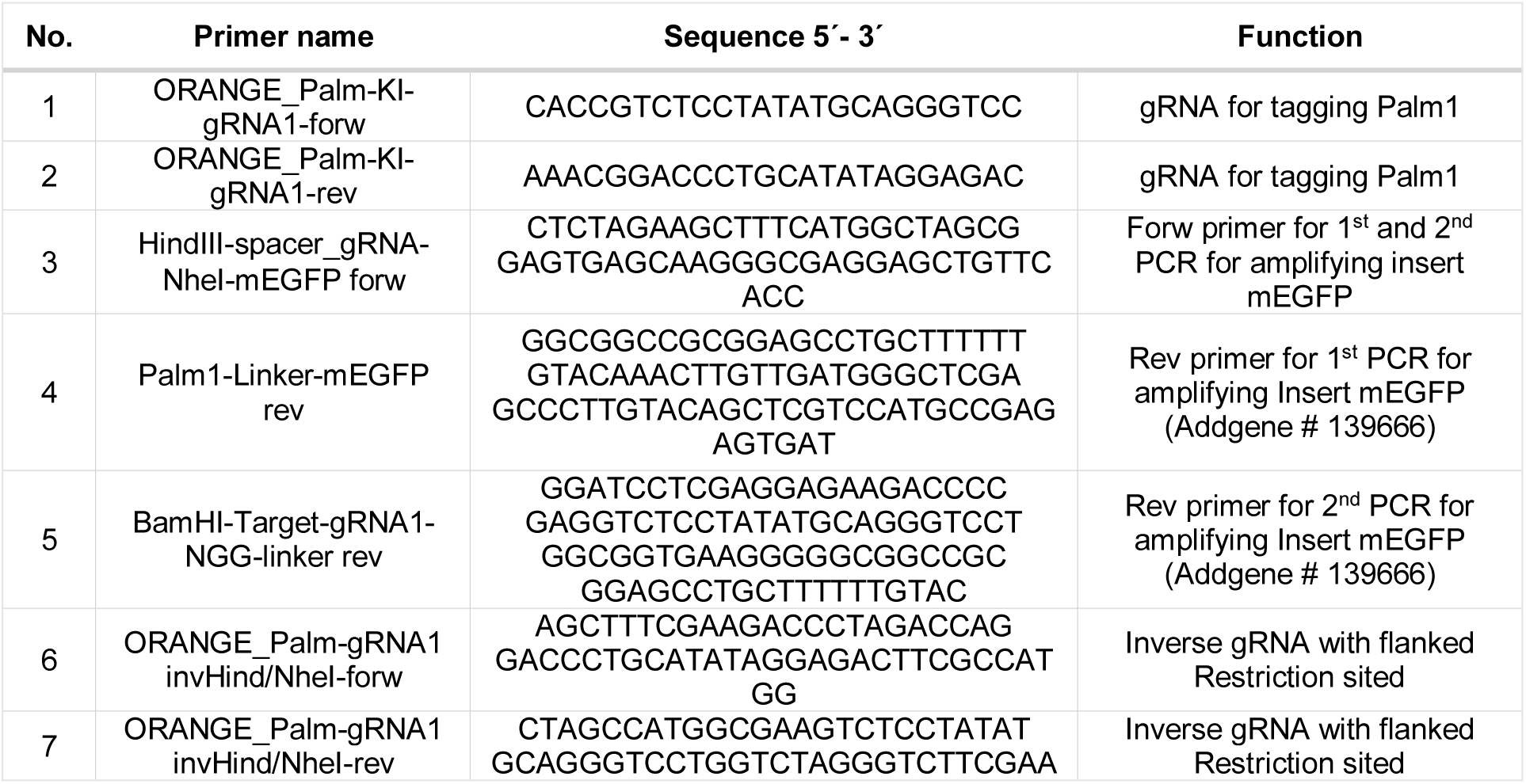
Oligonucleotides used for the generation of the mEGFP-CRISPR vector. Forw=forward; rev= reverse.

**Table 2:**
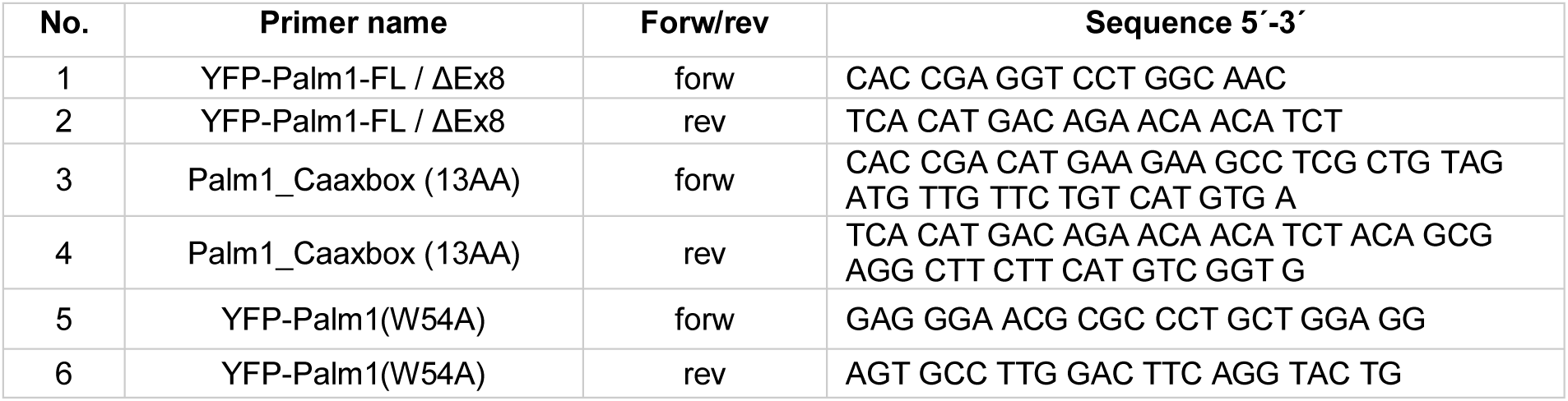
Oligonucleotides used for the generation of plasmids for YFP -Palm1 overexpression (primers 1-4) and W54A mutagenesis (primers 5-6). Forw=forward; rev= reverse.

**Table 3:**
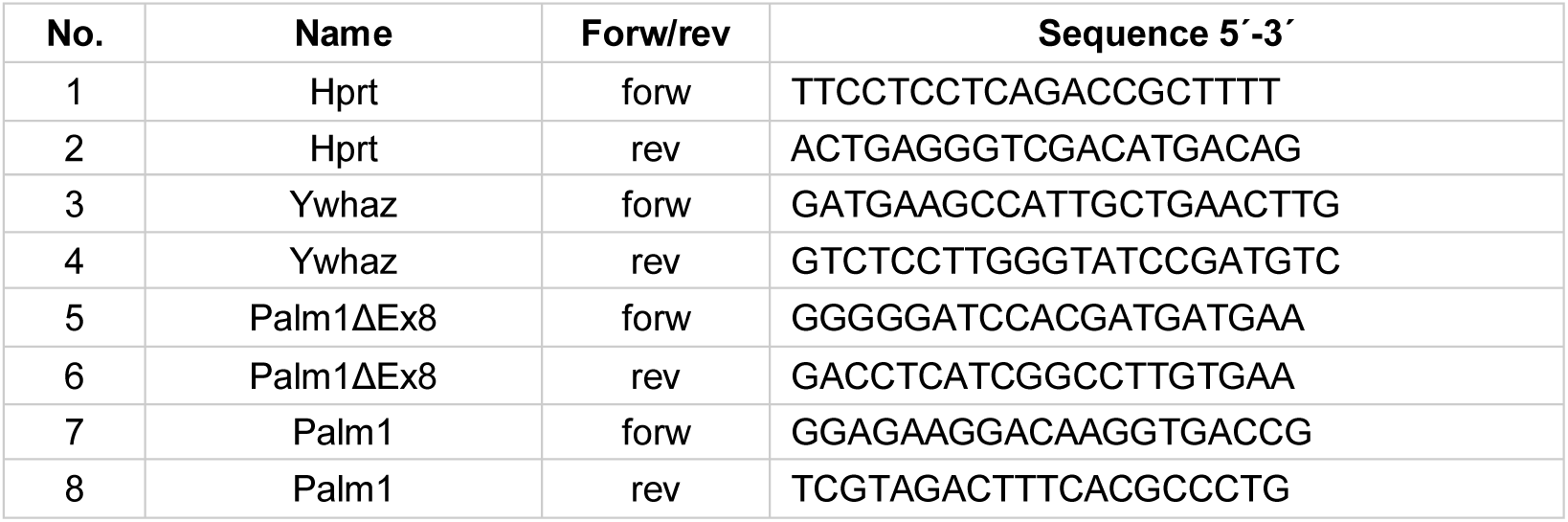
Oligonucleotides used for RT-qPCR experiments.

**Table 4:** 3D MINFLUX imaging sequence with an octahedral pattern.

| Iteration | Modality | L (nm) | Photon limit | Background limit | Pattern dwell time (ms) | Pattern repeat | Centre Frequency Ratio limit | Laser power factor |
| --- | --- | --- | --- | --- | --- | --- | --- | --- |
| 0 | Confocal | - | 160 | 15000 | 1 | 1 | - | 1 |
| 1 | z-line | - | 400 | 15000 | 1 | 1 | - | 1 |
| 2 | Square | 288 | 100 | 10000 | 1 | 5 | 0.8 | 1 |
| 3 | z-line | 288 | 50 | 10000 | 1 | 5 | - | 1 |
| 4 | Square | 151 | 67 | 10000 | 1 | 5 | - | 2 |
| 5 | z-line | 151 | 33 | 10000 | 1 | 5 | - | 2 |
| 6 | Square | 76 | 67 | 10000 | 1 | 5 | 0.8 | 4 |
| 7 | z-line | 76 | 33 | 10000 | 1 | 5 | - | 4 |
| 8 | Square | 40 | 100 | 10000 | 1 | 5 | - | 6 |
| 9 | z-line | 40 | 50 | 10000 | 1 | 5 | - | 6 |

### Yeast-2-Hybrid (Y2H)

The design of the Y2H bait for cDNA library screening was guided by the consideration that Palm1 is attached to the plasma membrane by a C-terminal lipid anchor (CaaX motif), while the N-terminus presumably reaches into the cytosol to engage other proteins^17^. Therefore, we chose the bait vector pFBL23^51^ in which the LexA DNA-binding domain (DBD) is fused to the Palm1 C-terminus after deletion of the CaaX box, leaving the N-terminus of Palm1 more accessible to interaction partners^52^. Palm1 and Palm1ΔEx8 cDNAs (without the last 13 codons; reference cDNA sequences Y14770 and Y14771) were generated by RT-PCR from mouse brain RNA, and were ligated into the EcoRI site of the vector pFBL23^51^ after blunting the EcoRI-cut ends with Klenow enzyme. The fusion proteins expressed under control of the ADH1 promoter encode mouse Palm1, beginning with its natural start codon, followed by a vector-encoded linker sequence of 9 aa and the LexA DNA-binding domain (DBD; layout of the expressed fusion protein: “N-Palm1-LexA-C”). This resulted in the following sequence context (vector sequences in lower-case letters, Palm1 and LexA-DBD coding sequences in upper-case letters): gaa ttg gag ATG GAG GTC…Palm1…GAT CTT gaa ttc gga tcc gga gtc gac ggt ggt ATG AAA GCG…LexA….

All Y2H assays were carried out at Hybrigenics (Evry, France). An adult mouse brain cDNA library in the prey vector pP6 (Hybrigenics), a derivative of pGADGH,^52^ was screened with the Palm1 (aa 1-370) bait construct, leading to the selection of 258 interaction-positive clones encoding βII-spectrin from 86 million library clones (Fig. 1A). The βII-spectrin interaction fulfilled Hybrigenics’ technical score for “very high confidence”, whereas no other interactors with “high” or “good confidence” were picked up in the screen. The screen was carried out at the unusually high concentration of 50 mM 3-amino- 1,2,4-triazol (3AT) to select for strongly interacting clones. 3AT is a competitive inhibitor in the histidine biosynthesis pathway that enhances the stringency of growth selection and gives a measure of the relative strength of bait-prey interaction. This explains why the screen produced many βII-spectrin clones but not a single clone representing the βI-, βIII- or βIV-spectrin isoforms, whose interactions with Palm1 are suppressed by 1-10 mM 3AT (Fig. 6E-F).

For further analysis of the Palm1-binding sequence features of βII-spectrin (Fig. 6D), an array of mouse βII-spectrin sequences (Spnb2 reference sequences: NP_787030.2, NM_175836.2) was generated by gene synthesis and cloned into prey vector pP7 (Hybrigenics) at Twist Bioscience (South San Francisco, USA), for subsequent 1-by-1 testing with the Palm1(1-370) bait. Bait and prey constructs were transformed in the yeast haploid cells L40ΔGal4 (mata) and YHGX13 (Y187ade2-101::loxP-kanMX-loxP, matα), respectively. The diploid yeast cells were obtained using a mating protocol with both yeast strains.^53^ These assays are based on the HIS3 reporter gene (growth assay without histidine). As negative controls, the bait plasmids were tested in the presence of empty prey vector (pP7) and all prey plasmids were tested with the empty bait vector (pFBL23), to confirm the absence of auto-activation. The interaction between Smad and Smurf was used as a positive control^54^ (Fig. 6F). Controls and interactions were tested by spotting aliquots of liquid cultures on DO-2 and DO-3 selective media. The DO-2 selective medium, lacking tryptophan and leucine, was used as a growth control, and to verify the presence of the bait and prey plasmids, in all Y2H experiments. The DO-3 selective medium, without tryptophan, leucine and histidine, selected for the interaction between bait and prey as shown in Fig. 6. To provide a semi-quantitative measure of the strength of Y2H interactions, all yeast cultures of Fig. 6D-H were spotted on DO-3 plates with a 3AT concentration series of 0, 1, 2, 5, 10, 20, 50, 100 and 200 mM.

To test for possible effects of phosphomimetic mutations on the Palm1 – βII-spectrin interaction, ten serine or threonine codons which are phosphorylated in mouse brain (S116, S124, T141, T145, T153, S157, T266, S345, T363, S365) (N.G.M. and M.W.K., unpublished) were replaced by phosphomimetic glutamate codons in pFBL23-Palm1(1-370), generating bait construct Palm1E5. Mutant Palm1 bait variants for the experiments of Fig. 6G+H were synthesized and cloned by Twist Biosciences into pFBL23, in exactly the same sequence context as the screening bait described above. As candidate preys collinear with the βII-spectrin construct G (aa 185-411; MKTA…LALR), the following mouse sequences of other β-spectrin/actinin isoforms (Fig. 6E) were cloned into pP7 at Twist Biosciences: Spnb1 (aa 185-411; MKTA…LALR; RefSeq: EDL36452), Spnb3 (aa 188-414; MKTA…LALR; RefSeq: NP_067262), Spnb4 (aa 193-419, MKTA…AALR; RefSeq: EDL24189), Spnb5 (aa 154-380, RKTA…QAMQ; RefSeq: NP_001351867), Actn1 (aa 156-383, RKTA…EWLL, RefSeq NP_598917), Actn2 (aa 163-390, RKTA…EWLL, RefSeq: NP001163796), Actn4 (aa 176-403, RKTA…EWLL, RefSeq: NP_001347477). Yeasts were transfected, mated and tested for growth as described above.

### Palm1 antibodies

Antisera against mouse Palm1 (aa 15-242 without exon 8) were raised in two rabbits (animals #2 and #10) as described and validated before^17,39^. Palm1 affinity-purified antibody #2 was used at 1:50 dilution for all immunofluorescence (IF) experiments shown in this study. Crude serum #2 was used at 1:50,000 dilution for Western blot experiments, but was also applicable for IF at 1:5000 dilution (not shown here). The specificity of the antibodies was confirmed under the conditions used here, by IF of neuronal cultures (Fig. S5A), and Western blotting of brain subcellular fractions (Fig. S6A-B) from WT and Palm1-KO animals.

### mEGFP-Palm1 CRISPR knock-in

pORANGE-mEGFP-Palm1 was based on the CRISPR/Cas9 knock-in template vector pORANGE (gift from H. MacGillavry, Addgene plasmid #131471; RRID:Addgene_131471)^55^. Palm1 sequence from Rattus norvegicus (NC_005106.4) was used as a template. GuideRNAs (gRNAs) close to the 5’-end of *Palm1* exon 2 were designed with the CRISPR design tool of Benchling (https://benchling.com) based on scoring algorithms (Table S1, primers #1-2)^56,57^. As a result, mEGFP flanked by linker sequences was inserted into codon 3 of Palm1 (ME-Linker1-mEGFP-Linker2-LAXX). For vector construction, the donor sequence, which contains mEGFP flanked by two inverse gRNA sequences, was generated by PCR and inserted into the pORANGE construct by restriction cutting with HindIII and BamHI, dephosphorylation and ligation (Table S1, primers #3-5). Finally, 5’ inverse gRNAs (upstream of EGFP) were annealed and ligated into the HindIII and NheI digested second intermediate construct (Table S1, primers #6-7). After bacterial transformation with the ligated plasmid, colonies containing the correct construct were identified by sequencing the purified plasmid (GeneJET Endo-free Plasmid Maxiprep Kit, Thermo Fisher cat. #K0861), which was later employed for the transfection of hippocampal neurons. The correct insertion of the mEGFP into exon 2 of the rat Palm1 gene was verified by sequencing the extracted genomic DNA of primary hippocampal rat neurons, at least 7 days after electroporation. The annotated sequence of the mEGFP-Palm1 knock-in is given in file S3.

### Plasmids for transient expression of proteins in mammalian cells

Palm1 coding sequences without N-terminal start codons but with C-terminal stop codons were amplified by RT-PCR from mouse brain single-stranded cDNA, generating two amplimers (Palm1(2-383) and Palm1(2-383)ΔEx8) with the same primer pair (Table S2, #1-2). The insert for YFP-CaaX (the C-terminal 13 codons of Palm1 directly fused to YFP) was synthesized in full-length, with the oligonucleotides #3 and #4 (Table S2) being annealed before cloning. Inserts were cloned via the Gateway^®^ system into a destination vector with an N-terminal YFP-encoding sequence (Invitrogen Vivid Colors^®^ pcDNA6.2/N-YFP-DEST). The coding sequences of all recombinant plasmids were confirmed to be mutation-free by sequencing (reference cDNA sequence Y14771). The plasmid without Palm1 insert was used to express YFP as a control. For the generation of YFP-Palm1(W54A) plasmid, the Q5^®^ Site-Directed Mutagenesis Kit (E0554, NEB) was used following manufacturer’s instructions. A primer-pair annealing to the template with their 5’ ends back-to-back were designed (Table S2 #5 (forward) and #6 (reverse)). Here, the first primer harboured the mutated bases. Primers were amplified together with the YFP-Palm1(2-383) plasmid before simultaneous kinase, ligase, and DpnI treatment. The success of the W54A mutagenesis and the absence of other mutations was confirmed by whole plasmid sequencing. βII-spectrin-HA was a gift from Vann Bennett (Addgene plasmid #31070; RRID:Addgene_31070).

### Palm1 knockout mice

Constitutive paralemmin-1 knock-out (Palm1-KO; Palm1-/-) mice were generated by targeted deletion of exon 5, causing a reading frame-shift, by TaconicArtemis (Köln, Germany), and maintained in a C57BL/6N genetic background. The official allele designation of this Palm1-KO mouse mutant is Palm^tm1.2Kili^ (MGI:6883588). The strain has been deposited in the European Mouse Mutant Archive (EMMA) under ID # EM:14921. Palm1-KO mice were viable, healthy and fertile without obvious phenotypic manifestations. Gene construct design and comprehensive KO mouse phenotyping will be described in more detail elsewhere. Western blot analysis demonstrated that this is a null mutation, with Palm1 protein expression in brain undetectable. Animals were genotyped by genomic PCR with reverse primer Palm1-int5R (acagacaggcatagaagttgc) and forward primers Palm1-3/ex5F (accttggtgaacgctcagca; WT-amplimer 375 bp) and Palm1-int4F (ccctacaacgctaacacttcc; mutant amplimer 315 bp). Mice were bred and kept in the mouse facility of the Max Planck Institute for Multidisciplinary Sciences (City Campus). All regulations given in §4 Animal Welfare Law of the Federal Republic of Germany (§4 TierSchG, Tierschutzgesetz der Bundesrepublik Deutschland) were followed. No specific authorization or notification was required for the breeding of animals, or for their sacrificing for organ dissection.

### Preparation of Primary Cultures of Hippocampal Neurons

All experimental procedures were performed in accordance with the Animal Welfare Act of the Federal Republic of Germany (Tierschutzgesetz der Bundesrepublik Deutschland, TierSchG) and the Animal Welfare Laboratory Animal Regulations (Tierschutz-versuchstierverordnung). Experiments were supervised by Animal Welfare officers of the Max Planck Institute for Medical Research (MPImF) and of the Max Planck Institute for Multidisciplinary Sciences (MPINAT), conducted and documented according to the guidelines of the TierSchG (permit number assigned by the MPImF: MPI/T-35/18 and MPI/T-36/18). No specific authorization or notification was required for the procedures performed in this study.

Primary hippocampal neurons were prepared from P0-P2 postnatal wild-type Wistar rats (Janvier-Labs, Le Genest-Saint-Isle, France), C57BL/6N mice or Palm1-KO mice of either sex. Briefly, isolated hippocampi were first digested with 0.25% trypsin for 20 min at 37 °C. The reaction was stopped by adding 1x DMEM supplemented with 10% heat-inactivated FBS (Thermo Fisher cat. 12491015 and cat. 10082147, respectively). Hippocampi were rinsed three times with Hanks solution (Thermo Fisher, cat. 10012011) before being mechanically dissociated by pipetting up and down in Neurobasal (NB) medium (Thermo Fisher, cat. 21103049) supplemented with 1% GlutaMAX (Thermo Fisher, cat. 35050061), 1% penicillin/streptomycin (Thermo Fisher, cat. 15070063) and 2% B27 (Thermo Fisher, cat. 17504044) (from here on referred to as supplemented NB). Dissociated neurons were passed through a 40 µm cell strainer (FisherbrandTM, cat. 22363547) and finally seeded on glass coverslips pre-coated with 0.1 mg/ml poly-L-ornithine (Sigma-Aldrich, cat. P3655) and 1 µg/ml laminin (Corning, cat. 354232). 110,000 cells were plated on ∅ 18 mm coverslips (12-well plate), while 55,000 cells were plated on ∅ 12 mm glass coverslips (24-well plate). For the experiments, in which neurons were transfected, 150,000 cells were plated on ∅ 18 mm coverslips (12-well plate). 1-2 hours after seeding, medium was changed to fresh supplemented NB, and on the following day 5 µM cytosine β-D-arabinofuranoside (AraC) was added to the cultures. Cells were incubated at 37°C, 5% CO2 until use. Neurons which were not immediately plated were frozen directly after dissociation at a concentration of 3-4 million cells/ml. The cell suspension was diluted 1:1 in freezing buffer (supplemented NB, 10% heat-inactivated FBS with 20% DMSO). Cryotubes were placed into a freezing container and stored at -80°C overnight before being transferred into liquid nitrogen until further use. For thawing, cryotubes were taken from liquid nitrogen and placed directly at 37 °C. Upon thawing, cells were diluted 1:1 with supplemented NB before being transferred to fresh supplemented NB (end concentration 800,000 cells/ml). 400,000 cells were plated on ∅ 18 mm coverslips (12-well plate) to achieve an expected final density of ∼110,000 cells/coverslip. Finally, medium was changed to supplemented NB 1 hour after seeding.

### Electroporation and transfection of neurons

Electroporation was performed at DIV 0 before seeding the cells with the Neon Transfection System (Thermo Fisher, cat. MPK5000) and the Neon^TM^ Transfection System 10 µl Kits (Thermo Fisher, cat. MPK1025) following manufacturer’s instructions. Briefly, 150,000 freshly dissociated neurons were washed once with 1x PBS, and mixed with 200 ng plasmid and 10 µl Buffer R. Cells were electroporated with 3 pulses of 10 ms each at 1400 V and immediately plated on ∅ 18 mm glass coverslips pre-coated as described above (12-well plate). Cells were kept in an incubator at 37°C and 5% CO2. 24 hours after seeding, medium was changed to fresh supplemented NB medium and 5 µM AraC was added to the cultures. Cells were further incubated under the same conditions until use.

Lipofection was performed at DIV 5 with Lipofectamine 2000 according to the manufactureŕs instructions (Thermo Fisher cat. 11668019). Before transfection, 600 µl of growth media was harvested and stored as preconditioned medium. Thereafter, 2 µg of plasmid and 2.5 µl Lipofectamine 2000 were separately incubated with 50 µl Opti-MEM respectively (Thermo Fisher, cat. 31985062) for 5 min at room temperature. Both solutions were then combined and incubated for 15 min before adding the mixture to cultures. 1 hour after transfection, neurons were transferred to preconditioned medium, filled up to 1 ml with fresh supplemented NB, and kept in culture until DIV 19. Neurons transfected with plasmid encoding βII-spectrin at DIV 5 were fixed at DIV 12.

### Electrophysiological analysis of hippocampal neurons

Cultured hippocampal neurons derived from WT or Palm1-KO mice were recorded at DIV 16-20, using whole-cell patch clamp in either current- or voltage-clamp configuration. For current-clamp recordings, the following internal solution was used (in mM): 125 K-Gluconate, 20 KCl, 10 HEPES, 0.5 EGTA, 4 MgATP, 0.3 NaGTP, 10 Na-phosphocreatine, osmolarity 312 mOsmol, pH 7.2 adjusted with KOH. For voltage-clamp recordings, the following internal solution was used (in mM): 125 Cs-gluconate, 20 KCl, 4 MgATP, 10 Na-phosphocreatine, 0.3 NaGTP, 0.5 EGTA, 2 QX314, 10 HEPES, 312 mOsmol, pH 7.2. In all recordings, the following extracellular solution was used (in mM): 125 NaCl, 2.5 KCl, 25 NaHCO3, 0.4 ascorbic acid, 3 myo-inositol, 2 Na-pyruvate, 1.25 NaH2PO4, 2 CaCl2, 1 MgCl2, 25 D(+)- glucose, 315 mOsmol, pH 7.4. The extracellular solution was continuously oxygenated with 95% O2 and 5% CO2. On the day of recording, a coverslip containing neurons was placed in a RC-27 chamber (Sutter Instruments), mounted under BX51 upright microscope (Olympus), equipped with DIC and fluorescent capabilities. Neurons were maintained at 26±1°C using a dual TC344B temperature control system (Sutter Instruments). Cells were approached and patched under DIC, using 3-4 MΩ glass pipettes (WPI) pulled with a PC100 puller (Narishige, Japan). In all experiments a Multiclamp 700B amplifier (Axon instruments, Inc) controlled by Clampex 10.1 and Digidata 1440 digitizer (Molecular Devices, Inc) was used. Detection and analysis of voltage- and current-clamp recordings was done with Clampfit 10.1 or with custom-written macros in IgorPro 6.11.

For current clamp experiments, automatic bridge-balance was performed after achieving whole-cell current-clamp configuration. The membrane potential in all neurons was maintained at approximately -70 mV by injecting the appropriate feedback current into the cells. Current injections <50 pA were considered acceptable. Action potentials were triggered by current injection through the recording pipette (500 ms, 25 pA steps from -200 to +400 pA). For voltage clamp recordings, the membrane potential was clamped at -70 mV. Series resistance <10 MΩ were considered acceptable, and were not electronically compensated. Miniature excitatory currents (recorded in the presence of 0.5 µM TTX) were detected as downward deflections, and analysed offline using Clampfit 10.4 and custom-written macros in IgorPro (Wavemetric).

### Immunostaining of cultured neurons for confocal and STED experiments

Neurons were briefly rinsed once with 1x PBS before fixation in 100% methanol for 10 min at -20°C, or in 4% PFA in PBS for 20 min at room temperature. PFA-fixed samples were quenched in quenching buffer (PBS, 100 mM glycine, 100 mM ammonium chloride) and permeabilized for 5 min in 0.1% Triton X-100 in PBS. Regardless of the fixation, samples were blocked with 1% BSA in PBS for 1 hour, and incubated with primary antibodies for 1 hour at room temperature in a wet and dark chamber, with the exception of the samples incubated with the Paralemmin-1 affinity-purified serum #2 antibodies, which were incubated overnight. After washing the samples five times with 1x PBS, secondary antibodies were added for 1 hour at room temperature. Finally, samples were embedded in Mowiol® 4-88 (Merck, Sigma-Aldrich cat. 81381) mounting medium supplemented with 2.5% w/w DABCO 33-LV (Merck, Sigma-Aldrich cat. 290734), according to the CSH protocol (https://cshprotocols.cshlp.org/content/2006/1/pdb.rec10255).

The following primary antibodies were used for immunostaining in this study, in addition to the Palm1 antibodies described above: βII-spectrin mouse (BD bioscience, cat. 612563, 1:400), ankyrin G guinea-pig (Synaptic Systems, cat. 386 005, 1:400), ankyrin G rabbit (Synaptic Systems, cat. 386 003, 1:400), ankyrin B mouse (NeuroMab, cat. 73-145, 1:10), βIII-tubulin chicken (Synaptic Systems, cat. 302306), MAP2 rabbit (Synaptic Systems, cat. 188002, 1:500), α-adducin rabbit (Abcam, cat. ab51130, 1:200). In case of multicolor experiments with primary antibodies raised in rabbit and guinea-pig, sequential staining was performed (*i.e.* first primary antibody in rabbit and corresponding secondary, and then the antibody in guinea pig and corresponding secondary).

The following secondary antibodies, nanobodies, and phalloidin conjugates were used at 1:100 dilution unless otherwise indicated: goat anti-mouse STAR635 (Abberior, cat. ST635P-1001), goat anti-rabbit STAR635P (Abberior, cat. ST635P-1002), goat anti-mouse STAR580 (Abberior ST580-1001), goat anti-rabbit STAR580 (Abberior, cat. ST580-1002), goat anti-guinea pig Alexa Fluor 488 (Thermo Fisher, cat. A-11073), goat anti-chicken Alexa Fluor 488 (Thermo Fisher, cat, A-21467), goat anti-rabbit Alexa Fluor 405 (Thermo Fisher, cat. A31556), FluoTag X-4 anti-GFP (Nanotag Biotechnologies, cat. N0304-Ab635P-S, 1:200), FluoTag®-X2 anti-PSD-95 STAR580 (Nanotag Biotechnologies, cat. N3702-Ab580L, 1:150), phalloidin-STAR635 (Abberior, cat. 2-0205-002-5). All analyses were done in rat neurons unless stated otherwise.

### Confocal and STED imaging

Confocal and STED images were performed on an Abberior Expert Line Microscope (Abberior Instruments GmbH, Germany) built on a motorized inverted IX83 microscope (Olympus, Tokyo, Japan) and equipped with pulsed STED lines at 775 nm and 595 nm, RESOLFT lines at 488 nm and 405 nm, excitation lasers at 355 nm, 405 nm, 485 nm, 580 nm, and 640 nm, and spectral detection. Spectral detection was performed with avalanche photodiodes (APD) and detection windows were set to 650-725 nm, 600-630 nm, 505-540 nm, and 420-475 nm to detect STAR635P, STAR580, Alexa Fluor 488/YFP/eGFP and Alexa Fluor 405, respectively. Confocal images were acquired either with a 20x/0.4 NA oil immersion lens with pixel size of 200 nm and 5 z-stack of 400 nm each, or with a 100x/1.4 NA oil immersion lens with a pixel size of 150 nm. The STED donut was generated with spatial light modulators (SLMs). STED images were acquired with the 100x/1.4 NA lens with a pixel size of 30 nm, and pinhole of 80 µm (0.8 AU). Laser powers and dwell times were adjusted for the different experiments but kept consistent for the different conditions within the same experiment. STED images displayed in Figure S7C have been acquired with a second Abberior Expert Line Microscope, described in the following FRAP experiments paragraph.

### Image processing and analysis

Acquired images were visualized by Imspector (Abberior Instruments GmbH) and processed by FIJI ImageJ 1.52p (https://fiji.sc). Axonal regions were classified in proximal (=AIS, defined as ankyrinG (ankG) IF-positive), middle (up to 40 µm after the AIS), and distal (further than 40 µm after the end of the AIS). This analysis was performed only on cultures older than 3 DIVs, since at earlier stages the axon could not be unambiguously identified.

Auto- and cross-correlation analyses were performed in Matlab2018b. Briefly, regions of interest of 1.5-2 µm long were manually selected and examined for periodic patterns using the function “xcorr2”, as described previously^11^. Shown are the amplitudes from the autocorrelation curve calculated as the difference between the value at 190 nm and the average of the values at 95 and 285 nm, where the first two valleys are expected.

Local intensities were measured from the confocal images in FIJI drawing manually line profiles of 6 µm along the axons including the same regions used for the correlation analysis. Values were normalized to the average calculated from the respective control of each experimental round (either non-transfected cells or YFP-transfected neurons) in the proximal regions. Per axon, one proximal, middle and distal region was measured.

For the growth cone analysis, central and peripheral domains were manually segmented based on the phalloidin channel. Transitional zone and peripheral domain were not distinguished.

Sholl analysis was performed in confocal images of neurons at DIV 3 with the FIJI tool “Sholl Analysis (From Image)”. For this analysis, a single neuron was identified and the rest of the components of the image were manually removed. Next, a threshold was set and all structures smaller than 0.5 µm were removed using “analyze particles”. After a median filter with radius 1, images were skeletonized. The centroid of the soma was calculated and set as the centre of the concentric shells. The distance between each shell was 1 µm. Numerous concentric shells were created until the last one did not intersect with any region of the neuron.

Neuronal stages were visually categorized according to ^25^. Briefly, stage 1 consists of neurons surrounded exclusively by growing lamellipodia. In stage 2, the premature neurites emerge. In stage 3, one neurite is clearly longer than the others and acquired axonal properties. Somata were segmented manually and their areas were calculated in FIJI.

PSD-95 clusters were analysed by applying first a Gaussian blur filter with sigma of 1 before setting a threshold of 17-300 counts. Then, the area and mean intensities of all PSD-95 particles with a size bigger than 0.02 µm^2^, selected using “analyze particles”, were calculated.

For display, brightness was adjusted uniformly throughout the images. Unless stated in the figure caption, no further image processing was performed and images are displayed as raw data.

### FRAP experiments

Neurons electroporated with plasmids containing YFP-Palm1, YFP-Palm1ΔEx8, or YFP at DIV 0 were imaged at DIV 14-16 on an Abberior Expert Line Microscope (Abberior Instruments GmbH, Göttingen, Germany) built on a motorized inverted IX83 microscope (Olympus, Tokyo, Japan). The microscope was equipped with pulsed STED lasers at 655 nm and 775 nm shaped by phase plates, and with 520 nm, 561 nm, 640 nm, and multiphoton (Chameleon Vision II, Coherent, Santa Clara, USA) excitation lasers. Spectral detection was performed with 2 avalanche photodiodes (APD) in the spectral window 530-560 nm. Images were acquired with a 60x/1.42 UPLXAPO60XO oil immersion objective lens (Olympus). Before imaging, samples were transferred into live magnetic chambers (Chamlide, Live Cell Instruments), washed once with prewarmed ACSF solution and stained with anti-neurofascin mouse (NeuroMab, Antibodies Incorporated, cat. 75-172, 1:400) diluted in ACSF for 5 min at 37 °C to identify the axons. After three washing steps, samples were incubated with goat anti-mouse Alexa Fluor 594 (Thermo Fisher, cat A-11032) diluted in ASCF 1:100 for 30 seconds and washed three times before imaging. Straight axonal regions of 20 µm x 5 µm were selected and imaged with 17 frames at a frequency of 1 Hz with a pixel size of 110 nm, 10 µs dwell time and 3 line accumulations as a reference before photo-bleaching. Subsequently, sharing the center of the region, a 4 µm x 2.5 µm region located at the center of the previous region of interest was bleached with two frames using the maximum 520 nm confocal laser power and the 660 nm STED laser with a 80 nm pixel size, 10 µs dwell time and 5 line accumulations. Fluorescence recovery was detected by imaging the initial 20 µm x 5 µm region for over 400 seconds at a frequency of 0.5 Hz for the first 30 seconds, and 1 Hz for the rest of the measurement with the same parameters used initially.

Fluorescence recovery was measured using FIJI ImageJ 1.52p (https://fiji.sc). Briefly, images were first segmented creating a grid of 1 µm x 1 µm. Axonal mean intensity within the photo-bleaching area (4 µm) was divided by the mean intensity of the reference area (2 µm, grid 2 and 3 starting from the border of the image) at every frame and normalized to the first one. After photo-bleaching, the mean intensity value of the first frame of the bleached area was subtracted for all the following frames, thus setting the initial fluorescence recovery to zero, and normalized by dividing it to by the reference area. Shown is the average of the obtained curves with the SEM.

### MINFLUX sample preparation and imaging

MINFLUX localized fluorophores by using a pattern of light featuring an intensity minimum and thereby achieves single-digit nanometer precision^33^. To this aim, DNA-PAINT was applied to achieve the blinking of single molecules^34^. In this technique, fluorescently labeled DNA oligomers transiently bind to the target structure decorated with the complementary oligomer. Neurons were electroporated with the plasmid encoding YFP-Palm1, fixed in 4% PFA between DIV 14 and 18, blocked in 1% BSA as described above, and incubated with primary rabbit antibody against α-adducin (Abcam, cat. ab51130, 1:200) overnight at 4 °C. Samples were then labeled with MASSIVE-TAG-Q-anti-GFP from the DNA-PAINT KIT of Massive Photonic following manufacturer’s protocol. Briefly, samples were incubated with single-domain anti-YFP nanobodies and anti-rabbit coupled to a single DNA-PAINT site respectively for 1 hour at room temperature (diluted 1:200 in antibody incubation buffer), and washed three times with washing buffer (all buffers from the DNA-PAINT KIT). Then, samples were incubated with gold nanorods (Nanopartz Inc. A12-40-980) diluted 1:1 in 1x PBS or gold Colloid (BBI Solutions, #SKU EM.GC150/7) for 5 minutes at room temperature, and washed 3 times with 1x PBS. Samples were finally transferred into a magnetic chamber with ∼0.5 nM imager 3 anti-GFP-Atto 655 diluted in imaging buffer and imaged for at least 30 min. For exchange DNA-PAINT, after recording of the GFP channel, the medium was changed several times until no further valid localizations were detected. Finally, ∼1.5 nM imager 2 anti-rabbit-Atto 655 was added and the same region was imaged for adducin.. MINFLUX imaging was performed on an Abberior 3D MINFLUX (Abberior Instruments GmbH, Göttingen, Germany) built on a motorized inverted microscope IX83 (Olympus, Tokyo, Japan) and equipped with 640 nm, 561 nm, 485 nm, and 405 nm laser lines. Detection was performed with 2 APDs in the spectral windows 650-685 and 685-720. Images were acquired using the default 3D imaging sequence, with an L in the last iteration step of 40 nm, and photon limit of 100 and 50 photons for the lateral and axial localization, respectively (imaging sequence provided in Table S3). The localization precision attained in these experiments was: σx=7.2; σy=6.3; σz=7.6 nm with 50 photons per dimension.

### Analysis of MINFLUX data

Drift correction was performed as in ^35^. Briefly, localizations were divided into time windows and rendered as images to further calculate the 3D correlation function between different time points. To obtain the estimated 3D drift path, the cross-correlation between time windows was used. The final drift trajectory was then subtracted from the localization coordinates.

Localizations from the same emission trace, *i.e.* with same trace identification (TID), farther than three standard deviations with respect to the mean trace position were excluded from the trace. Only the traces containing at least 4 localizations were considered. The experimental localization precision was estimated by computing the median value of the trace standard deviation^58^. Drift-corrected and filtered localizations corresponding to axonal regions varying from 2 to 4 µm were manually brushed from the original region. The center axis (*i.e.* line that passes along the center of an axon) was estimated and a coordinate transform was applied to have the axis parallel to the x-coordinate. To calculate the central axis, the brushed points were projected on a third-order polynomial fit for XY projection and in first-order polynomial fit for XZ projection. The aligned structure was divided in sections of about 50 nm length along its main axis and the localizations of each section were projected in a transverse plane. The best fit for an ellipse according to Least Squares criterion was found for each section and expanded along the main axis to form an elliptical cylindrical shell. The process was iteratively repeated until the center of the fitted ellipses matched the center of the central axis. Finally, to create an unwrapped view, the localizations were mapped to the corresponding shell along the radial direction, resulting in the relative position of the localizations along the axon circumference. The standard Euclidian distance was used to calculate the nearest neighbor distances from the unwrapped positions of TIDs, which already accounts for the axon curvature.

### Subcellular fractionation and Western blotting

Subcellular fractionation was performed to investigate the protein levels in brains from juvenile (postnatal day 9) male C57BL/6N and Palm1-KO mice (five biological replicates, each), all procedures being performed on ice or at 4 °C. Each brain (∼300 mg) was homogenized in 1400 µl homogenization buffer, with 10 strokes at 900 rpm in a 2 ml glass/teflon piston homogenizer. Homogenization buffer: 320 mM sucrose, 3 mM MgCl2, 1 mm EDTA, 1 mM DTT, 20 mM Tris pH 7.5, plus protease and phosphatase inhibitors. The homogenates were spun in a microliter centrifuge for 10 min at 900g = 3000 rpm, resulting in fractions P1 (nuclei & debris) and S1. S1 was re-centrifuged for 20 min at 10,000g = 9500 rpm, yielding fractions P2 (putatively mitochondria and synaptosomes) and S2. S2 was re-spun for 30 min at 30,000g = 16,300 rpm, producing fractions P3 (putatively plasma membranes and cytoskeleton) and S3 (putatively light membranes [”microsomes”] and cytosol). Pellets were each resuspended in 100 µl homogenization buffer, and all fractions were aliquoted and flash-frozen in liquid nitrogen. Protein concentrations were determined by Bradford assay (Bio-Rad, cat. 5000202). 15 µg of each lysate were mixed with 4x Laemmli buffer (Bio-Rad, cat. 1610747) supplemented with 10% 2-mercaptoethanol at a 4:1 ratio (Merck, Sigma-Aldrich, cat. M6250) and heated at 55 °C for 7 min. Samples were resolved by SDS-PAGE on 4-15% gradient Mini-PROTEAN® TGX precast protein gel (Bio-Rad, cat. 4561086) in 1x Tris/glycine/SDS buffer (Bio-Rad, cat. 1610732) at 85 V for 100 minutes. Proteins were wet-transferred onto a 45 µm pore size Immun-Blot® LF PVDF membrane (Bio-Rad, cat. 162-0260) in methanol transfer buffer (25 mM Tris pH 7.5, 190 mM glycine and 20% methanol) at 120 V for 85 minutes. Membranes were blocked with 3% BSA/PBS for 1 hour before incubating the primary antibodies diluted in 1% BSA overnight at 4 °C under gentle rotation. The following day, membranes were rinsed three times for five minutes with 1x TBST (TBS, 0.1% Tween-20 (Carl Roth, cat. 9005-64-5)) and secondary antibodies diluted in 1% BSA were incubated for 1 hour at room temperature under gentle rotation. Finally, membranes were rinsed three times for five minutes with 1x TBST and visualized with the UVP ChemStudio PLUS (Analytikjena). Precision Plus Protein Dual Color Standards (Bio-Rad, cat. 1610374) were used as reference.

Two proteins were detected simultaneously, followed by a stripping step of 30 min at 50°C under gentle rotation (stripping buffer: 6.25% (v/v) 1M Tris pH 6.7, 10% (v/v) 20% SDS, 0.7% (v/v) β-mercaptoethanol) and a blocking step as described above. Order of detection was: (i) βII-spectrin mouse (BD bioscience, cat. 612563, 1:1500) with goat anti-mouse Alexa Fluor 488 (Thermo Fisher, cat. A32727), and Palm1 crude serum rabbit (1:50,000) with goat anti-rabbit Alexa Fluor 647 (Thermo Fisher, cat. A-21245), (ii) Kv1.2 mouse (NeuroMab, cat 73-008, 1:50) with goat anti-mouse Alexa Fluor 488, and Tomm20 rabbit (Abcam ab186735, 1:1500) with goat anti-rabbit Alexa Fluor 647, (iii) GAPDH rabbit (Cell Signalling, cat. #2118, 1:1500) with goat anti-rabbit Alexa Fluor 647. Dilution of secondary antibodies 1:1500. Finally, the signal of βII-spectrin and Palm1 of each fraction were normalized to their respective average signal from the WT-P2 fraction.

### RNA extraction and reverse transcription quantitative real-time PCR (RT-qPCR)

RNA extraction and RT-qPCR were performed as described previously^59^. Briefly, frozen rat primary neurons (DIV 3, 7, 14 or 20) from 4 - 5 ∅ 18mm coverslips were combined to create 1 biological replicate. Total RNA was extracted using RNeasy Mini Kit (Qiagen, cat. 74104) following manufacturer’s protocol. Briefly, cells were disrupted within the wells and homogenized using Lysis buffer RTL, and transferred to QIAshredder spin columns (Qiagen, cat. 79656). Genomic DNA was removed by adding RNase-free DNase I (Qiagen, cat. 79254) for 15 min at room temperature. RNA was eluted with 30 µl of RNase free water. RNA quality and concentration were measured by UV spectrophotometry on NanoDrop One (Thermo Fisher, cat. ND-ONE-W). RNA samples with absorbance ratios below 1.0 were precipitated using ammonium acetate, glycogen (Thermo Fisher, cat. R0551) and 100% ethanol following manufacturer’s protocol. The High-Capacity RNA-to-cDNA kit (Thermo Fisher, cat. 4387406) was used to reverse-transcribe RNA to cDNA.

RT-qPCR was performed using PowerUp SYBR Green Master Mix (Thermo Fisher, cat. A25742), with 40 ng cDNA and 20 µM of each primer in a 10 µl reaction volume per sample. All RT-qPCR experiments were performed in biological triplicate on a LightCycler 480 (Roche) and gene expression levels were normalized to the expression of the housekeeping genes Hprt and Ywhaz, using oligonucleotides in Table S3. Non-baseline-corrected RT-qPCR raw data were extracted from the LightCycler 480 software to provide input for LinRegPCR (version 2020.0), and analyzed accordingly to ^59^. Briefly, the software performed baseline correction for each sample individually and calculated amplification efficiency (E), quantification cycle (Cq) and coefficient of determination (R2) by fitting a linear regression model to log-linear phase. Technical replicates were examined and arithmetic mean was taken as the Cq value representing biological samples.

### Statistical analysis and preparation of figures

Statistical tests and the plotting of graphs were performed with Origin 2019 or GraphPad Prism8. Statistical tests are described in each caption. Correlation coefficients were interpreted as suggested in ^60^: 0.0-0.3 no correlation, 0.3-0.5 low correlation, 0.5-0.7 moderate correlation, 0.7-1 high correlation. Statistical differences are indicated as follows: * = p<0.05; ** = p<0.005, *** = p<0.005.

The drawing of Fig. 8 was created with Biorender.com with the “Academic License Rights”. Colors in panel (I) of figure 6 were modified using PyMOL™ 2.5.5. All figures were assembled in Adobe Illustrator 2022.

## Acknowledgements

We thank Prof. Stefan W. Hell and Prof. Nils Brose for supporting this project. We acknowledge Christian Kutzleb (formerly with M.W.K. at the University of Bochum) for constructing the Palm1 and Palm1ΔEx8 bait vectors in pFBL23, and Kathrin Kusch and Christiane Senger-Freitag (Institute for Auditory Neurosciences, Göttingen) for re-cloning the Palm1E5 insert into pFBL23, and Petra Tafelmeyer (Hybrigenics) for expert support. We thank Francisco Balzarotti for developing the MATLAB script to measure the periodicity. We are grateful to Katrin Willig for access to laboratory spaces; Jana Kress, Birgit Koch, Alena Fischer, and Valerie Dürr for support with neuronal cultures; Daniel Bollack for fruitful discussions. Gabriele Paolo Malengo for advice on FRET experiments; and the staffs of the animal facility and the genotyping facility of the MPI for Multidisciplinary Sciences (City Campus) for Palm1-KO mouse maintenance, breeding, and genotyping. This research was funded by the Deutsche Forschungsgemeinschaft (DFG, SFB1286/A07 to E.D.; and grant KI 324/14 to M.W.K), and the Swedish Research Council (Vetenskapsradet 2003-3398 to M.W.K.).

## Author contributions

V.M.P. performed and analyzed all imaging experiments and Western blots, supervised by E.D.; M.T. performed and analyzed qRT-PCR experiments, supervised by A.P.; C.A. performed and analyzed electrophysiology experiments; J.H. designed the CRISPR/Cas9 construct and provided technical support with neuronal cultures; J.H., N.G.M., and S.K. developed and validated reagents; M.A.d.R.B.F.L. analysed MINFLUX data; V.M.P. prepared figures; M.W.K. initiated the study by suggesting a possible involvement of Palm1 in the MPS to E.D., designed and supervised the Y2H analyses, and provided the Palm1-KO mice,; V.M.P., E.D. and M.W.K. conceived the experiments and conceptualized the project; V.M.P., E.D., and M.W.K. wrote the manuscript with input from all authors.

