## Supplementary Information for "Paralemmin-1 controls the nanoarchitecture of the neuronal submembrane cytoskeleton"

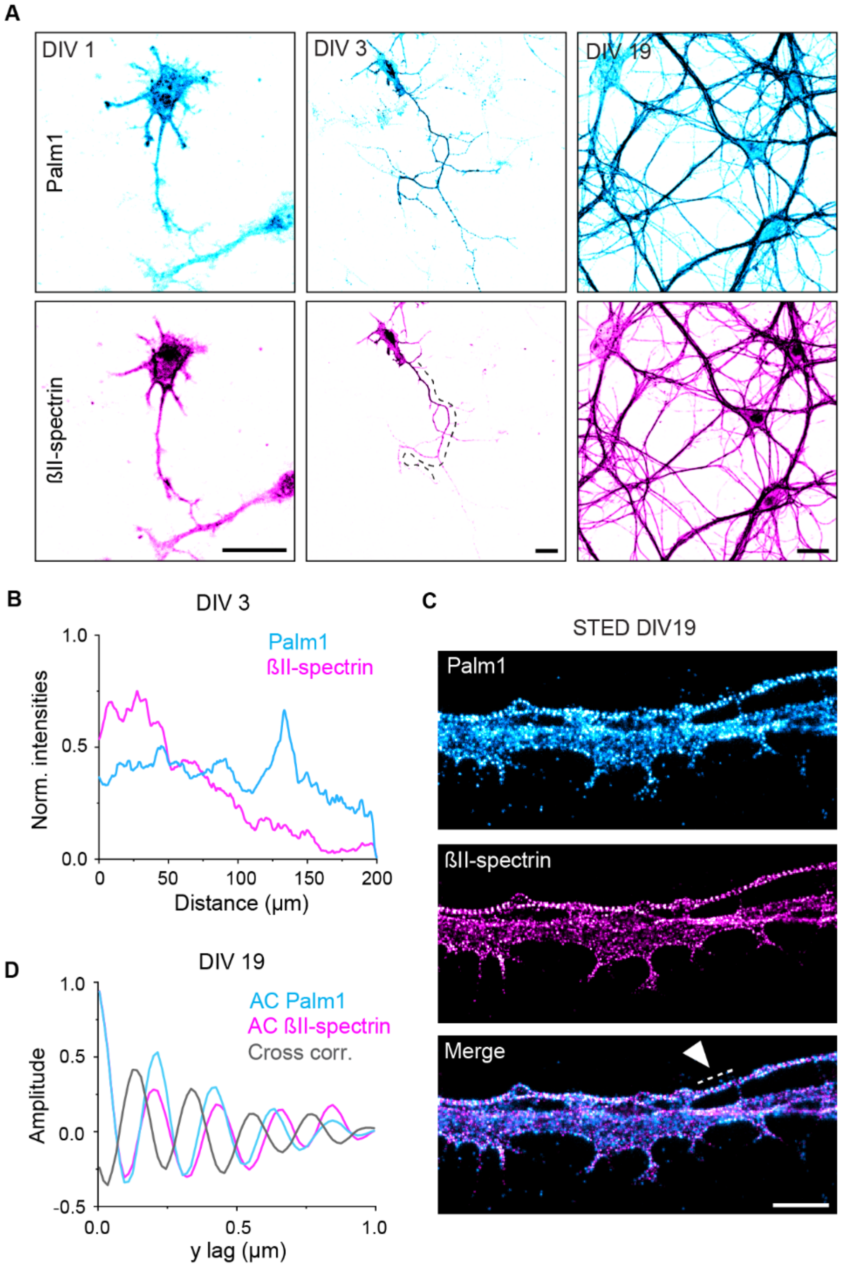


***Figure S1: The cellular and nanoscale organization of Palm1 is conserved between rats and mice.*** *(A) Representative confocal images of mouse-derived HPNs at DIV 1, 3 and 19, immunolabeled against Palm1 and ßII-spectrin (methanol fixation). (B) Normalized line profile of intensities (A.U.) and smoothed (50 values) for the axon indicated by the dashed line in (A) shows that Palm1 is present in distal regions before ßII-spectrin. (C) Two-color STED image of an axon and a dendrite shows a clear periodic pattern of Palm1, intercalating with ßII-spectrin, especially along the axon. Scale bar: 2 µm. (D) Autocorrelation (AC) and cross-correlation (CC) analyses performed along the axon, in the region indicated by the dashed line in (C).*

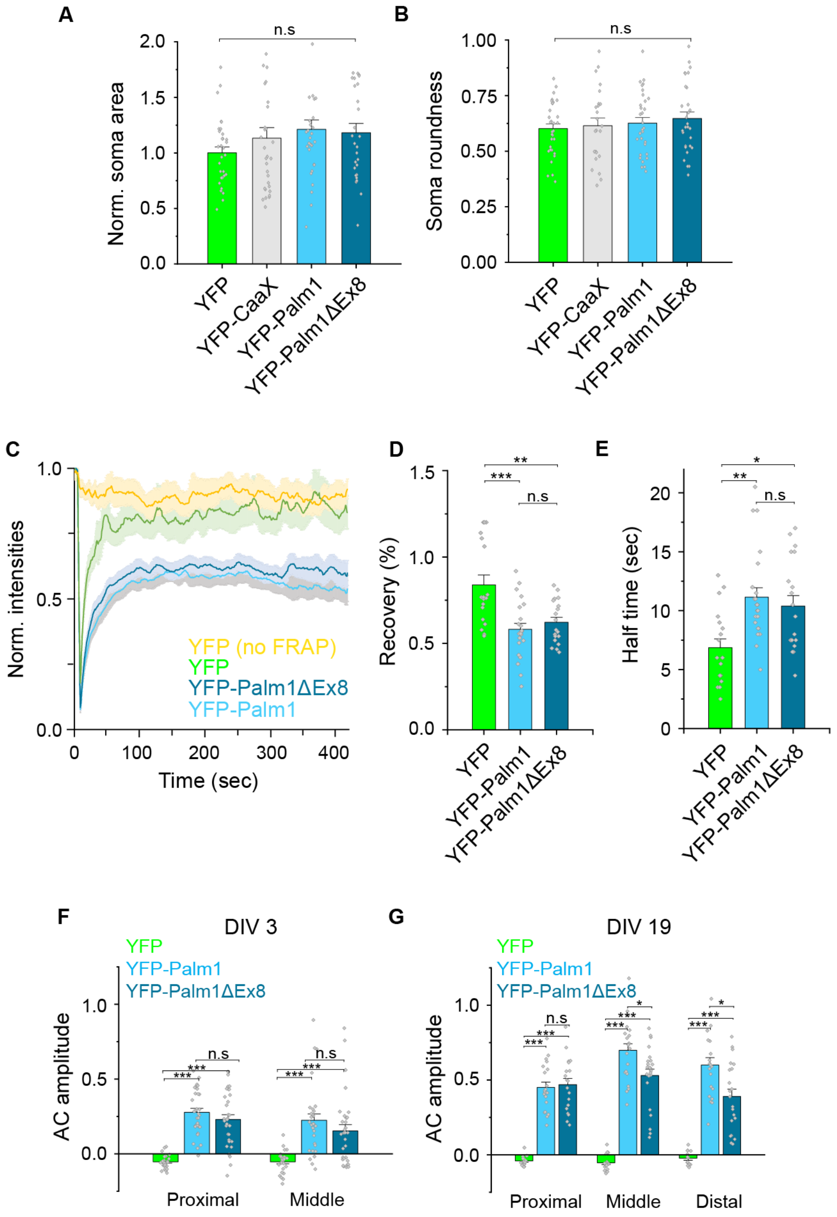


***Figure S2: Characterization of Palm1 splice variants upon overexpression.*** *(A) Normalized soma area (A.U.) and (B) roundness of rat HPN at DIV 3 are unaffected by overexpression of YPF, YFP-CaaX, YFP-Palm1, or YFP-Palm1ΔEx8. Cells analyzed: YFP, n=34; YFP-CaaX, n=24; Palm1, n=31; Palm1ΔEx8, n=27. All from N=3 independent neuronal cultures.* ***(C-E) Palm1 splice variants exhibit similar diffusion properties in mature neurons (DIV 14-16).*** *(C) FRAP analysis of mature neurons overexpressing Palm1 or Palm1ΔEx8 show a reduced fluorescence recovery (D) and half-time (E) compared to control neurons overexpressing YFP. Solid lines indicate mean values while shades indicate the standard error. Axons analyzed: YFP-Palm1 n=24, YFP-Palm1ΔEx8 n=20, YFP n=18. All from N=3.* ***(F-G) Recombinant Palm1 exhibits an increased periodic organization along the proximal, middle and distal axon upon overexpression****. AC amplitudes of Palm1 and Palm1ΔEx8 along axons at DIV 3 (F) and DIV 19 (G), detected with a nanobody against YFP. Axons analyzed in the proximal/middle region (DIV 3): YFP-Palm1, 31/30; YFP-Palm1ΔEx8, 31/31. All from N=3. Axons analyzed in the proximal/middle/distal region (DIV 19): YFP-Palm1, 20/21/20; YFP-Palm1ΔEx8, 19/23/21. All from N=3. Statistical analyses: One-way ANOVA; p-values in file S2.*

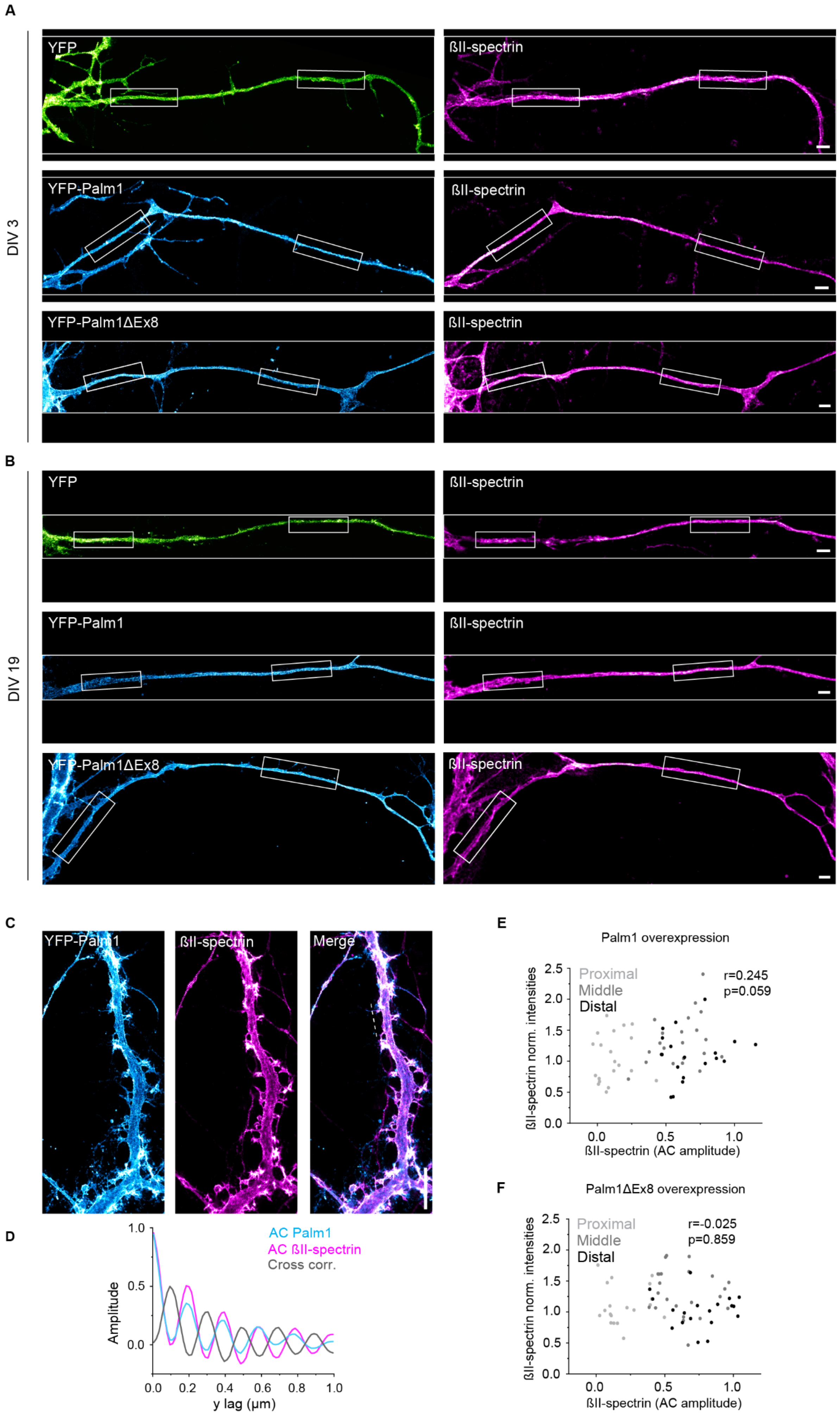
A
DIV 3
YFP
ßII-spectrin
YFP-Palm1
ßII-spectrin
YFP-Palm1ΔEx8
ßII-spectrin
B
DIV 19
YFP
ßII-spectrin
YFP-Palm1
ßII-spectrin
YFP-Palm1ΔEx8
ßII-spectrin
C
YFP-Palm1
ßII-spectrin
Merge
D
Amplitude
AC Palm1
AC ßII-spectrin
Cross corr.
y lag (µm)
E
Palm1 overexpression
Proximal
Middle
Distal
r=0.245
p=0.059
ßII-spectrin norm. intensities
ßII-spectrin (AC amplitude)
F
Palm1ΔEx8 overexpression
Proximal
Middle
Distal
r=-0.025
p=0.859
ßII-spectrin norm. intensities
ßII-spectrin (AC amplitude)

***Figure S3: (A-B) Entire axons from which the proximal and middle regions shown in Fig. 3 marked with the white box have been cropped.*** *(A) Representative STED images of axons of neurons overexpressing YFP (green), YFP-Palm1 (cyan) or YFP-Palm1ΔEx8 (cyan), immunolabeled against YFP and ßII-spectrin (magenta) at DIV 3 (A) and (B) DIV 19 (B). Scale bars: 2 µm.* ***(C-F) Palm1 overexpression enhances ßII-spectrin periodicity also in dendrites.*** *(C) Representative STED image of a dendrite of mature neurons (DIV 19, PFA fixation) overexpressing YFP-Palm1 and immunostained against YFP and endogenous ßII-spectrin. Scale bar: 2 µm. (D) AC and CC analyses of the dendritic region marked by the dashed line in the merged image in (C) confirm the clear and long-range, alternating periodic organization of both proteins. (E-F) Correlation scatter plot of the local intensities vs. periodicities of ßII-spectrin after the overexpression of Palm1 (E) or Palm1ΔEx8 (F) display no correlation.*

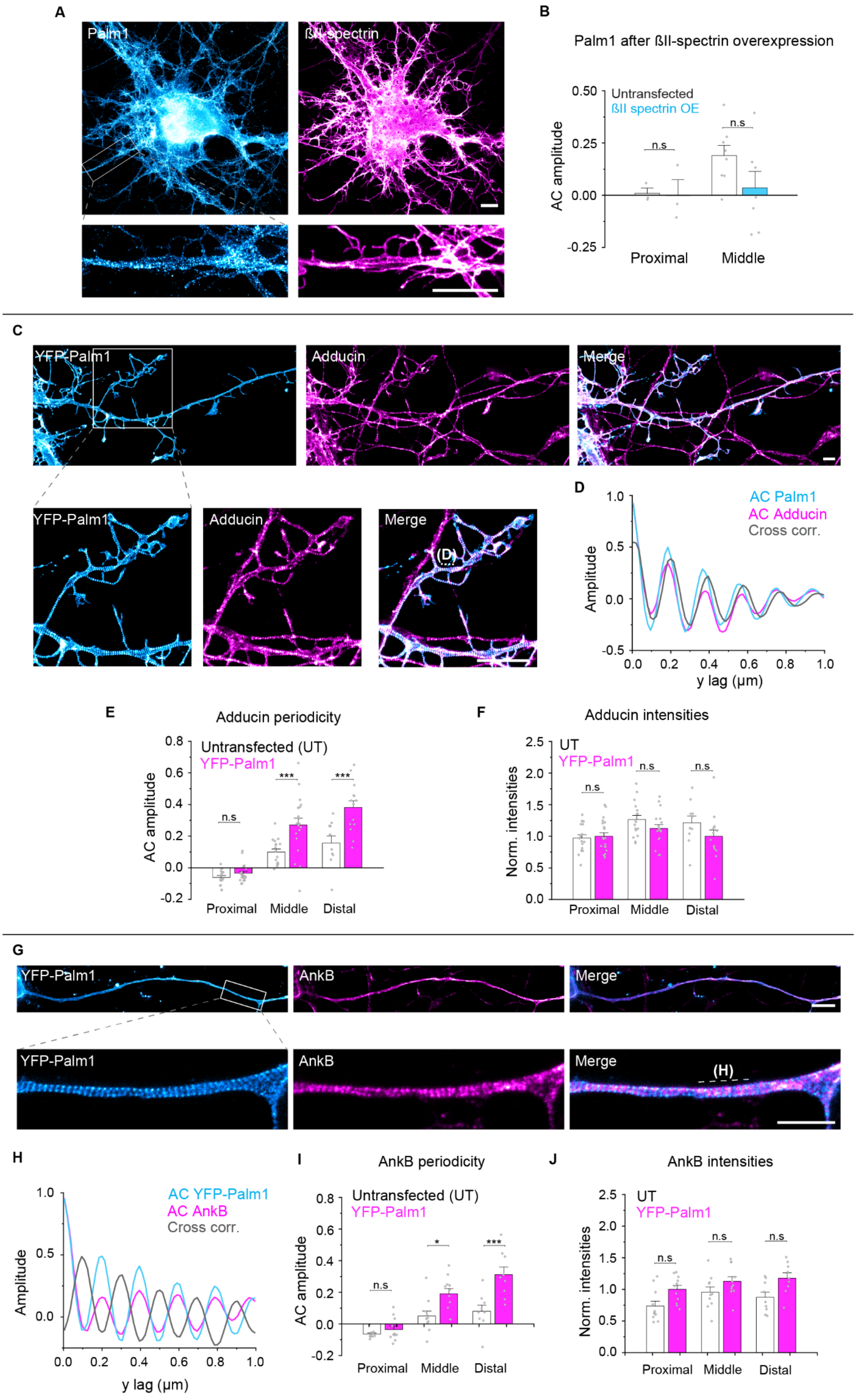
A
Palm1
ßII-spectrin
B
Palm1 after ßII-spectrin overexpression
Untransfected
ßII spectrin OE
AC amplitude
0.50
0.25
0.00
-0.25
n.s
n.s
Proximal
Middle
C
YFP-Palm1
Adducin
Merge
YFP-Palm1
Adducin
Merge
(D)
D
AC Palm1
AC Adducin
Cross corr.
Amplitude
1.0
0.5
0.0
-0.5
0.0
0.2
0.4
0.6
0.8
1.0
y lag (µm)
E
Adducin periodicity
Untransfected (UT)
YFP-Palm1
AC amplitude
0.8
0.6
0.4
0.2
0.0
-0.2
n.s
***
***
Proximal
Middle
Distal
F
Adducin intensities
UT
YFP-Palm1
Norm. intensities
2.5
2.0
1.5
1.0
0.5
0.0
n.s
n.s
n.s
Proximal
Middle
Distal
G
YFP-Palm1
AnkB
Merge
YFP-Palm1
AnkB
Merge
(H)
H
AC YFP-Palm1
AC AnkB
Cross corr.
Amplitude
1.0
0.5
0.0
0.0
0.2
0.4
0.6
0.8
1.0
y lag (µm)
I
AnkB periodicity
Untransfected (UT)
YFP-Palm1
AC amplitude
0.8
0.6
0.4
0.2
0.0
-0.2
n.s
*
***
Proximal
Middle
Distal
J
AnkB intensities
UT
YFP-Palm1
Norm. intensities
2.5
2.0
1.5
1.0
0.5
0.0
n.s
n.s
n.s
Proximal
Middle
Distal

***Figure S4: (A-B) Overexpression of ßII-spectrin does not affect the nanoscale organization of Palm1.*** *(A) Representative STED image of a neuron transfected at DIV 5 and PFA-fixed at DIV 12 overexpressing ßII-spectrin, immunostained for endogenous Palm1 and ßII-spectrin. Scale bar: 5 µm. (B) AC amplitudes of Palm1 in the proximal and middle axon in untransfected neurons and upon ßII-spectrin overexpression. Number of axons analyzed in the proximal/middle axon: WT: 3/8; ßII-spectrin overexpression: 3/7 from N=3.* ***(C-J) Palm1 overexpression enhances the periodic organization of adducin and ankB in the middle and distal axon without altering their local concentrations.*** *(C) Representative STED images showing the nanoscale organization of overexpressed YFP-Palm1 and endogenous adducin in a mature neuron at DIV 19, transfected at DIV 5 (PFA fixation). White box indicates region shown in the close-ups below. Scale bar: 2 µm. (D) Representative AC and CC analyses of both proteins performed along the region indicated by the dashed line in the merged image. (E) AC amplitude and (F) normalized intensities (A.U.) of adducin in untransfected neurons conditions and upon Palm1 overexpression. Axons analyzed in the proximal/middle/distal region: WT: 17/17/10; Palm1 overexpression: 21/19/14. All from N=3. (G) Representative STED images showing the nanoscale organization of overexpressed YFP-Palm1 and endogenous ankB along the axon of a mature neuron at DIV 19, transfected at DIV 5 (PFA fixation). White box indicates axonal region shown in the close-ups below. Scale bars: 2 µm. (H) Representative AC and CC analysis of both proteins performed along the region indicated by the dashed line in the merged image. (I) AC amplitude and (J) normalized intensities (A.U.) of ankB in untransfected conditions and after overexpression of Palm1. Axons analyzed in the proximal/middle/distal region: WT: 11/11/9; Palm1 overexpression: 11/11/9. All from N=3. All statistical analyses in B-J by one-way ANOVA; p-values in file S2.*

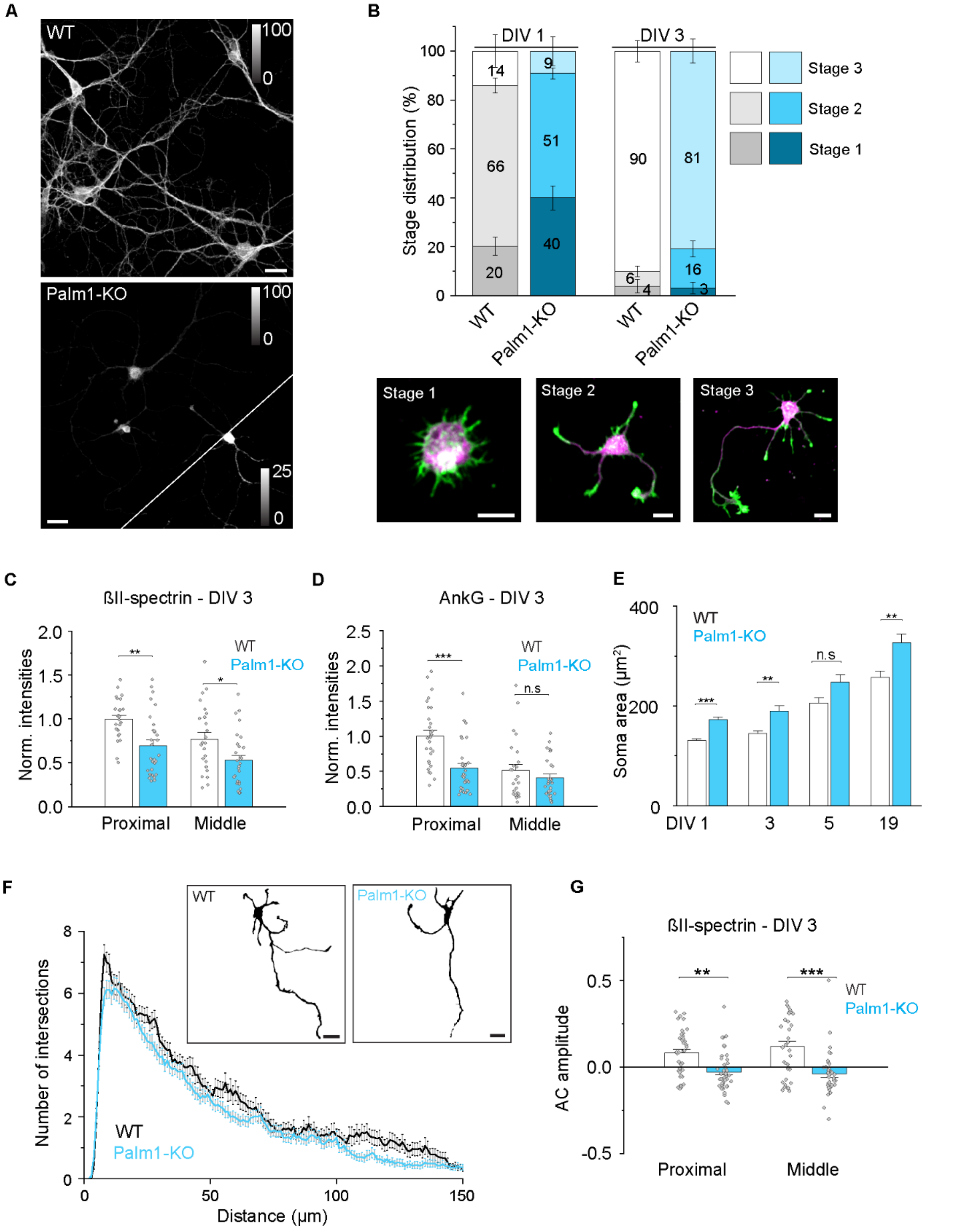


***Figure S5: Palm1-KO delays neuronal development****. (A) The depletion of Palm1 was confirmed by Palm1-IF in Palm1-KO neurons fixed in PFA at DIV 12. Representative confocal images of WT (upper) and Palm1-KO (lower) hippocampal neurons. Lower right corner of the Palm1-KO image is shown with enhanced brightness, as indicated by the look-up table values. Scale bars: 25 µm. (B) Stage distribution of WT and Palm1-KO hippocampal neurons at DIV 1 and DIV 3 according to Dotti et al. 1988. Representative examples from Palm1-KO HPN are shown below, green: phalloidin, magenta: ßII-spectrin. Scale bars: 10 µm. Number of cells analyzed WT/Palm1-KO: DIV 1, 475/437; DIV 3, 181/162. (C) Normalized intensities (A.U.) of ßII-spectrin and (D) ankG along the proximal and middle axons of neurons at DIV 3. Number of WT/Palm1-KO axons analyzed: Proximal, 27/30; Middle, 25/28. All from N=3. (E) Soma area of WT and Palm1-KO neurons during in vitro maturation. Number of WT/Palm1-KO somata analyzed: DIV 1, 402/388; DIV 3, 192/168; DIV 5, 102/125; DIV 19, 61/91. All from N=3. (F)*

*Sholl analysis of WT and Palm1-KO neurons at DIV 3 showed a slightly reduced branching, illustrated by the representative confocal images. Scale bar 20 µm. (G)* ***The periodic organization of ßII-spectrin is affected by the Palm1-KO already early in development****. Number of WT/Palm1-KO axons analyzed: Proximal, 38/42; Middle, 36/39. All from N=3. Statistical analyses: One-way ANOVA; p-values in file S2.*

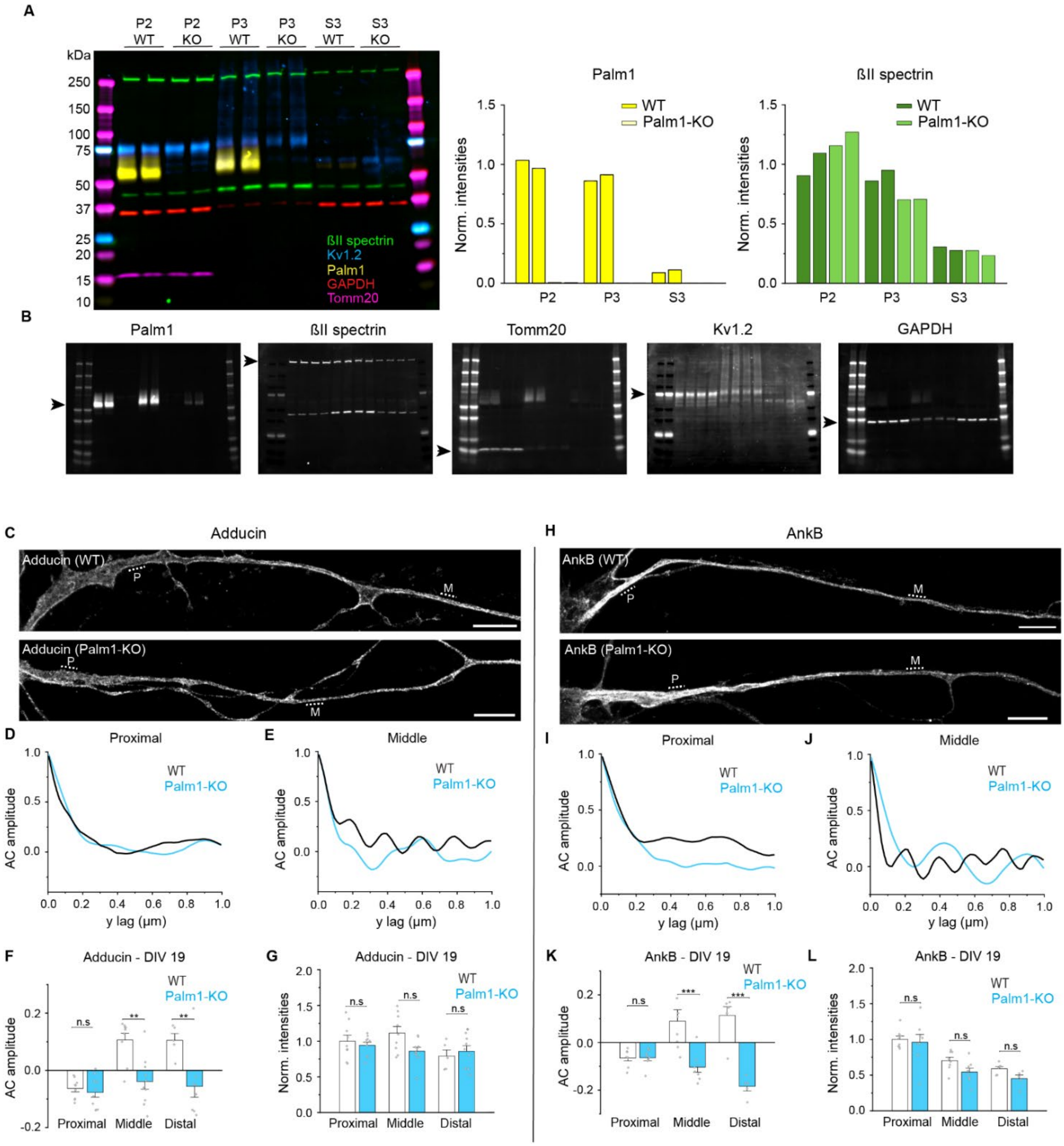

***Figure S6: ßII-spectrin concentrations in subcellular fractions of juvenile (P9) mouse brain are unaffected by constitutive Palm1-KO, as demonstrated by Western blot analysis.*** *(A) (Left) Merged image of a 5-color Western blot developed for Palm1, ßII-spectrin, Tomm20 (marker for mitochondria), Kv1.2 (marker for plasma membranes), and GAPDH (marker for cytosol). (Right) Quantification of the intensities obtained for Palm1 and ßII-spectrin in the different fractions, normalized to the averaged intensities calculated for P2-WT for each protein. (B) Individual Western blot channels. Black arrows indicate the expected bands. Precision Plus Protein Standards (Bio-Rad) were used. P2: mitochondria/synaptosomes, P3: plasma membrane/cytoskeleton, S3: microsomes/cytosol. Fractions from 2 WT and 2 Palm1-KO animals were analysed.* ***(C-G) Palm1-KO abolishes adducin and ankB periodicity in mature hippocampal neurons.*** *(C) Representative STED images of adducin along axons of WT and Palm1-KO neurons (DIV 19). Scale bars: 5 µm. (D) Representative AC amplitudes of the periodic organization of adducin along the proximal and (E) middle axon regions highlighted in (C)*

*by the dashed lines (P: proximal; M: middle) (F) AC amplitudes of the periodic organization of adducin and (G) local intensities measured along the same axonal regions on which the AC was calculated of WT and Palm1-KO neurons. Axons analyzed for AC amplitudes WT/Palm1-KO: Proximal: 9/10; Middle: 9/10; Distal: 5/9. All from N=1. (H-L) Same as in (C-G), but for ankB. Axons analyzed: Proximal: 8/8; Middle: 8/8; Distal: 6/5. All from N=1. One-way ANOVA; p-values in file S2.*

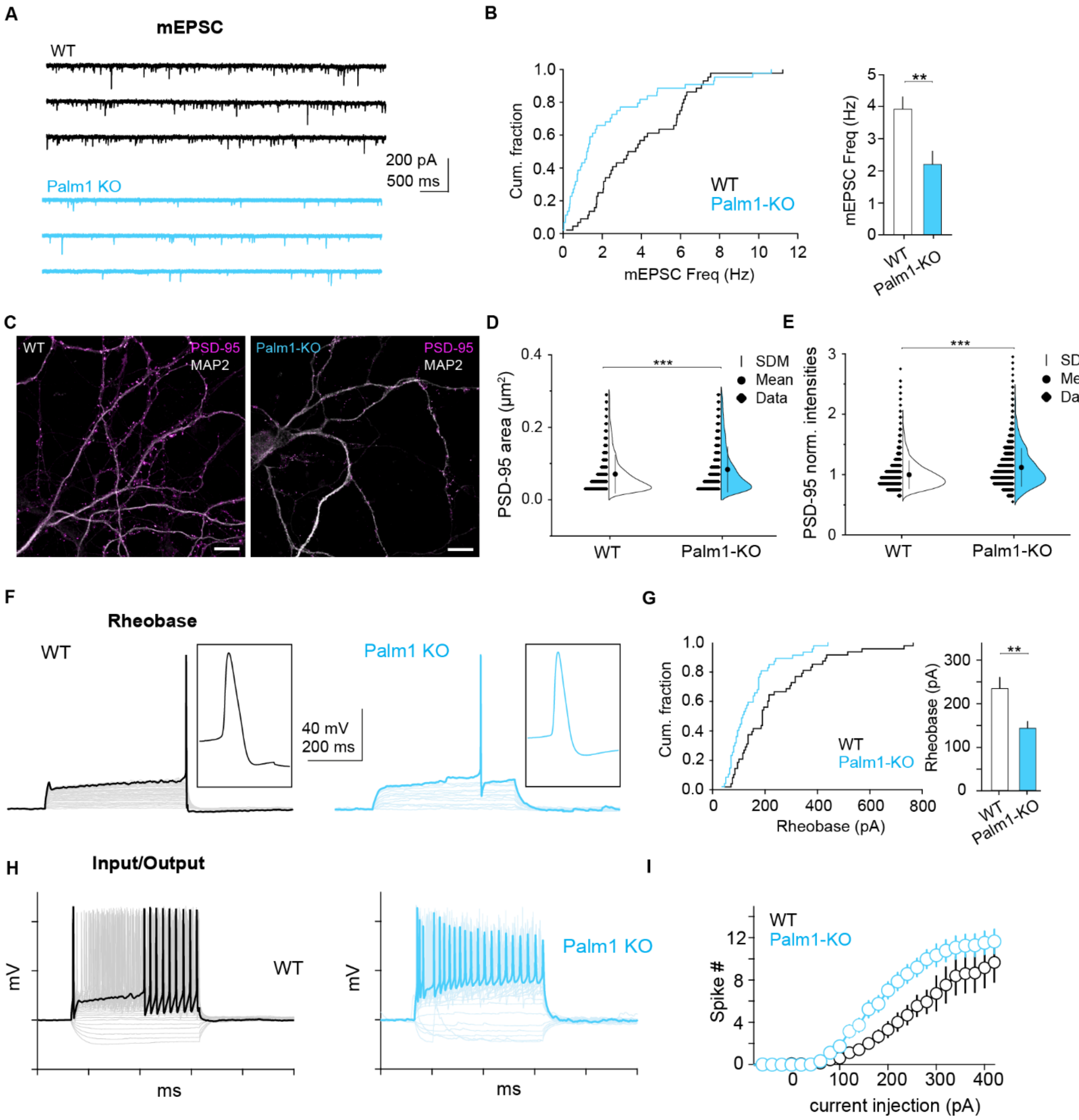


***Figure S7: Palm1-KO affects electrophysiological and postsynaptic parameters of hippocampal neurons.*** *(A) Representative miniature EPSC (mEPSC) traces recorded at -70 mV from mature (DIV 16-22) WT (black) or Palm1-KO (blue) neurons. (B) Left: Cumulative fraction of mEPSC frequency in WT and Palm1- KO neurons. Right: Plot of the averaged mEPSC frequency (mean±SEM) in Palm1-WT and KO neurons. Cells analyzed: WT: 44, Palm1-KO: 44. All from N=4. Unpaired t-test with Welch´s correction; p-value=0.002. (C) Palm1-KO results in fewer PSD-95 clusters but with a more intense fluorescent signal, as shown in the representative STED images of WT (left) and Palm1-KO (right) hippocampal neurons immunostained for PSD-95 and MAP2. Scale bar: 10 µm. (D) Area and (E) normalized fluorescence intensity of PSD-95 clusters in neurons lacking Palm1 compared to WT neurons. Mann-Whitney test; p-value for (D): $4.5x10^{-14}$. p-value for (E): $8.95x10^{-62}$. Number of PSD-95 clusters analyzed: WT/Palm1-KO: 3982/2855 from N=2 independent neuronal cultures. (F) Representative rheobase measurements (5 pA steps for 500 ms) in a WT and a Palm1-KO neuron. Evoked spikes are enlarged on the right. (G) (left) Cumulative distribution and (right) averaged (mean±SEM) rheobase measured in WT and Palm1-KO neurons. Palm1 depletion affects the rheobase, requiring less voltage for the generation of action potentials. Cells analyzed: WT: 48; Palm2-KO: 47. All from N=4. Unpaired t-test with Welch´s correction; p-value=0.002. (H) Representative voltage responses of a WT and a Palm1-KO neuron upon current injections of increasing amplitude (25*

*pA steps for 500 ms from -100 pA to 400 pA). The responses to 200 pA current injection are highlighted in bold. (I) Summary plot of the spike # as a function of current injected in WT and Palm1-KO neurons. Each data point corresponds to the mean±SEM. Number of cells analyzed: n=48 for WT and n=46 for Palm1-KO conditions. All from N=4.*

***Video S1: Animation of the N-terminal region of βII-spectrin.*** *The sequences relevant for the interaction with Palm1 are color-coded as in Fig. 6I.*

***Video S2****:* ***Movie of the 3D MINFLUX image shown in figure 7E****. First, exclusively Palm1 (cyan) is visualized being periodically localized at the plasma membrane of an axon. Thus, it allows the navigation along the resulted hollow structure. Performing exchange DNA-PAINT, the periodic pattern of adducin (magenta), being flanked by Palm1 (cyan), is observed. Scale bar is shown in figure 7E.*
